# Nucleus Accumbens Fast-Spiking Interneurons Constrain Impulsive Action

**DOI:** 10.1101/516609

**Authors:** Marc T. Pisansky, Emilia M. Lefevre, Cassandra L. Retzlaff, Brian H. Trieu, David W. Leipold, Patrick E. Rothwell

**Author notes:** Corresponding Author: Patrick E. Rothwell, Ph.D., 4-283 Wallin Medical Biosciences Building, 2101 6^th^ Street SE, Minneapolis, MN, 55455.

## Abstract

**Background:** The nucleus accumbens (NAc) controls multiple facets of impulsivity, but is a heterogeneous brain region with diverse microcircuitry. Prior literature links impulsive behavior in rodents to gamma aminobutyric acid (GABA) signaling in the NAc. Here, we studied the regulation of impulsive behavior by fast-spiking interneurons (FSIs), a strong source of GABA-mediated synaptic inhibition in the NAc.

**Methods:** Male and female transgenic mice expressing Cre recombinase in FSIs allowed us to identify these sparsely distributed cells in the NAc. We used a 5-choice serial reaction time task (5-CSRTT) to measure both impulsive action and sustained attention. During the 5-CSRTT, we monitored FSI activity with fiber photometry calcium imaging, and manipulated FSI activity with chemogenetic and optogenetic methodology. We used electrophysiology, optogenetics, and fluorescent *in situ* hybridization to confirm these methods were robust and specific to FSIs.

**Results:** In mice performing the 5-CSRTT, NAc FSIs showed sustained activity on trials ending with correct responses, but declined over time on trials ending with premature responses. The number of premature responses increased significantly after sustained chemogenetic inhibition or temporally delimited optogenetic inhibition of NAc FSIs, without any changes in response latencies or general locomotor activity.

**Conclusions:** These experiments provide strong evidence that NAc FSIs constrain impulsive actions, most likely through GABA-mediated synaptic inhibition of medium spiny projection neurons. Our findings may provide insight into the pathophysiology of disorders associated with impulsivity, and inform the development of circuit-based therapeutic interventions.

## INTRODUCTION

Impulsivity is defined as the tendency to act prematurely or without foresight, and involves a failure of cognitive control that characterizes various neuropsychiatric diseases (1, 2). Notably, impulsivity represents a vulnerability marker for substance use disorders (3), attention deficit/hyperactivity disorder (ADHD) (4, 5), and suicidality (6). The nucleus accumbens (NAc), a major hub in limbic circuits that regulate reward-seeking behavior, plays a critical role in controlling multiple facets of impulsivity (7). Evidence for this role comes from human brain imaging studies (8–11), as well as local pharmacological manipulations and brain lesion studies in rodents (12–17). These approaches have identified key neuromodulatory systems and provided critical insight into the aggregate function of the NAc, but we still have a limited understanding of how impulsive behavior is regulated by the activity of specific cell types embedded within NAc microcircuitry.

Gamma aminobutyric acid (GABA) has been implicated as a modulator of impulsivity in the NAc (18, 19) and other brain regions (20). A major source of local GABAergic signaling in these brain regions are fast-spiking interneurons (FSIs), a cell type frequently defined by parvalbumin (PV) expression (21). FSIs provide robust feed-forward inhibition that constrains and sculpts the output of projection neurons in the NAc (22–25), dorsal striatum (26–28), and other brain regions (29–31). FSIs in the prefrontal cortex regulate complex cognitive processes like attention (32), while FSIs in the dorsal striatum have been implicated in learning (33, 34) and habit formation (35). Manipulations of FSIs in the NAc alter behavioral responses to addictive drugs (36, 37), but the specific contribution of these cells to impulsivity has yet to be explored.

In this study, we investigated the function of NAc FSIs in mice performing the 5-choice serial reaction time task (5-CSRTT), a classic behavioral assay that measures sustained attention and impulse control (38). We focused on FSIs in the NAc core subregion, where GABAergic signaling has specifically been linked to impulsive behavior in this task (18). To monitor the activity of FSIs in behaving mice, we used a viral approach to express a genetically-encoded calcium indicator, and monitored fluorescent signals using *in vivo* fiber photometry. These calcium imaging experiments reveal that sustained activity of FSIs is associated with successful control of impulsive action. We then used chemogenetic and optogenetic methods to inhibit the activity of FSIs in the NAc core, and found these manipulations increased impulsivity. Together, our data suggest that FSIs in the NAc play an important role in constraining impulsive actions.

## METHODS AND MATERIALS

### Subjects

Male and female mice C57/B6J mice were housed with same-sex littermates on a 12 hr light/dark cycle, and used for experiments at 8-20 weeks of age. PV-2A-Cre transgenic mice (39) were obtained from The Jackson Laboratory (JAX stock #012358). All procedures conformed to the National Institutes of Health *Guidelines for the Care and Use of Laboratory Animals*, and were approved by the University of Minnesota Institutional Animal Care and Use Committee

### Behavioral Training

As previously described (40), mice were food-restricted to ~85% of free-feeding body weight, and pre-exposed to 14 mg purified dustless precision pellets (Bio-Serv) in the home cage. Behavioral training took place in standard mouse operant chambers (Med Associates), and began with two consecutive days of magazine training (30 food pellets delivered randomly over 30 minutes). We then began the first of eight training stages (Table 1), using a protocol adapted from previous studies of rats (41) and mice (42, 43). For additional details, see Supplemental Methods.

**Table 1.** Training protocol for the 5-CSRTT.

| Training Stage | ITI Length (sec) | Cue Duration (sec) |
| --- | --- | --- |
| 1 | N/A | ∞ (all cues) |
| 2 | N/A | ∞ (single cue) |
| 3 | 2.5 | 20 |
| 4 | 5 | 10 |
| 5 | 5 | 8 |
| 6 | 5 | 4 |
| 7 | 5 – 10 | 4 |
| 8 | 5 – 10 | 2 |
| Impulsivity Test | 2.5, 5, 10, 20 | 5 |
| Attention Test | 5 | 2.5, 5, 10, 20 |

### Stereotaxic Surgery

Intracranial virus injection and fiber-optic implantation were performed as previously described (40), with minor modifications described in Supplemental Methods. Viral vectors are described in the Key Resources Table and were used at a concentration of 3-5×10^12^ particles/mL. After surgery, mice were given 500 µL saline and 5 mg/kg carprofen (s.c.) daily for three days, and recovered a minimum of 7 days before any behavioral testing.

### Fiber Photometry

Fiber photometry recordings were conducted a minimum of two weeks after surgery, to allow for sufficient viral expression. Recordings from mice in the 5-CSRTT were conducted after one day of training in Stage 8. Real-time data were acquired as previously described (44), with details provided in Supplemental Methods.

### Chemogenetic Manipulations of Behavior

PV-2A-Cre mice and wild-type littermates (negative control group) received bilateral injection of AAV8-hSyn-FLEX-hM4Di-mCherry into the NAc. After recovery, mice began training on the 5-CSRTT through Stage 8, providing at least four weeks of viral expression. After one day of training in Stage 8, mice received i.p. injections of either saline or 2 mg/kg CNO (Hello Bio) 30 minutes prior to testing. Saline and CNO sessions were conducted on separate days in counterbalanced order. After these tests, mice completed attention and impulsivity tests, again preceded by counterbalanced i.p. injections of saline or CNO (2 mg/kg).

### Optogenetic Manipulations of Behavior

Separate groups of PV-2A-Cre mice received bilateral injections of either AAV5-EF1a-DIO-eNpHR3.0-mCherry or AAVdj-EF1a-FLEX-eYFP (negative control group) into the NAc, and bilateral implantation of 200um, 0.6NA fiber optic implants positioned +0.2mm dorsally. Mice were then trained on the 5-CSRTT through Stage 8, providing at least three weeks of viral expression prior to testing. Upon reaching Stage 8, mice were habituated to fiber optic tethering until achieving >50% correct. Optogenetic stimulation (593nm, constant power) was delivered using a laser (Shanghai Dream Lasers), coupled to a two-channel fiber-optic commutator, and controlled by TTL signals from the behavioral control software. Continuous light power was limited to ~3mW to minimize tissue heating (45). Mice were tested with light delivery in Stage 8 as well as impulsivity and attention tests.

### Histology

We confirmed the location of fiber-optic implants and extent of viral infection with immunohistochemistry, as described in Supplemental Methods. One animal from the optogenetic experiment was excluded due to lack of eNpHR3.0 viral expression.

### Fluorescent *in situ* Hybridization

Fluorescent *in situ* hybridization was performed using the RNAscope Multiplex Fluorescent Assay (Advanced Cell Diagnostics), as described in Supplemental Methods.

### Statistics

Analysis of variance (ANOVA) was conducted in IBM SPSS Statistics v24, with details provided in Supplemental Methods. All summary data are displayed as mean +/− SEM, with individual data points from male and female mice shown as closed and open circles, respectively. Complete statistical results can be found in **Table S1**.

## RESULTS

### Behavioral Measures of Impulsivity and Attention in Female and Male Mice

The 5-CSRTT requires mice to withhold a nose-poke response until it can be directed to one of five locations indicated by a brief visual cue (Figure 1A). Correct responses to the illuminated nose-poke aperture were rewarded with a food pellet, while all other trial outcomes were punished with a time-out period. These other outcomes included premature responses during the inter-trial interval (ITI) before cue presentation, incorrect responses to the wrong location, and omission trials in which the mouse fails to make a timely response. Training consisted of eight sequential stages of increasing ITI and decreasing cue durations (Table 1). To examine behavioral performance under varying degrees of task difficulty, we systematically varied the duration of either the visual cue (to test attention) or the ITI (to test impulsivity), as previously described (43). In a pilot study, we conducted these attention and impulsivity tests both early and late in training (Figure 1B). In the attention test, omission responses increased monotonically with briefer cues (F_1.59,22.26_=86.54, p<0.001), and decreased overall from early to late training (F_1,14_=6.11, p=0.027; Figure 1C). In the impulsivity test, premature responses increased monotonically as the ITI grew longer (F_1.74,24.35_=67.93, p<0.001), and decreased overall from early to late training (F_1,14_=22.21, p<0.001; Figure 1D).

**Figure 1.**
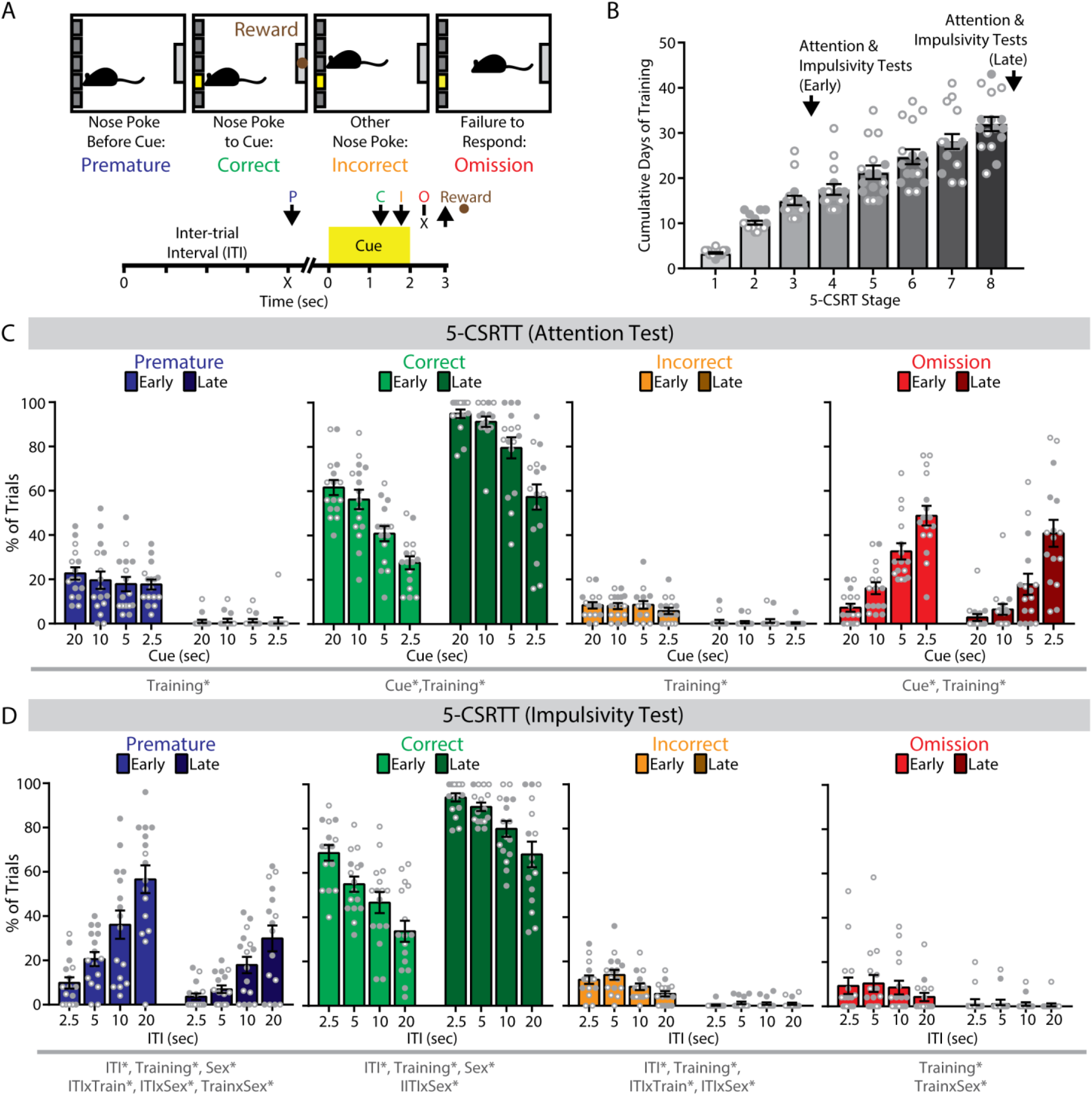
The 5-choice serial reaction time task (5-CSRTT) in mice. (**A**) (**top**) Trial outcomes in the 5-CSRTT. (**bottom**) Timeline of individual trials in the 5-CSRTT. Arrows indicate nose poke response. P, premature; C, correct; I, incorrect; O, omission. (**B**) Progression of mice in 5-CSRTT training. (**C**) In a test of attention at early and late training phases of the 5-CSRTT, percentage of correct, and omission trials varied by cue duration. (**D**) In a test of impulsivity at early and late training phases of the 5-CSRTT, the percentage of premature, correct, and incorrect trials varied by inter-trial interval (ITI) duration. n = 16 mice (8 female, 8 male).*p<0.05 for the indicated main effect or interaction.

This cohort included equal numbers of female and male mice. Total days required to complete the final stage of training was similar between sexes (F_1,14_=1.58, p=0.23), but there was a significant Stage x Sex interaction (F_7,98_=2.80, p=0.011). Female mice completed stages 2 and 7 more rapidly, while male mice completed stages 1 and 3 more rapidly (p<0.05, LSD post-hoc test). In the impulsivity test, there were significant Training x Sex x ITI interactions for correct responses (F_2.83,39.63_=3.55, p=0.025) and premature responses (F_2.90,40.54_=5.14, p=0.005), as well as a significant Training x Sex interaction for omissions (F_1,14_=8.28, p=0.012). Early in training, female mice had more omissions (F_1,14_=6.24, p=0.026), while male mice made more premature responses (F_2.71,37.98_=13.42, p<0.001) and fewer correct responses (F_2.72,38.09_=8.81, p<0.001) as the ITI grew longer. However, these sex differences in task performance were no longer detected late in training (all F<1). We used this protocol in subsequent experiments to investigate how NAc FSIs regulate 5-CSRTT performance in well-trained mice of both sexes.

### Characterization of Optogenetically-Evoked Calcium Signals from FSIs

To study FSIs in the NAc core, we used a mouse line expressing Cre recombinase in a bicistronic fashion from the *Pvalb* locus (PV-2A-Cre). In this mouse line, Cre is active in cells with both high and low *Pvalb* expression (39), providing robust labeling of FSIs in both the dorsal striatum (33, 35) and NAc shell (22, 36). Stereotaxic injection of Cre-dependent adeno-associated virus (AAV) labeled cells with electrophysiological characteristics of FSIs (**Figure S1A-C**), including a high maximum firing rate, narrow action potential half-width, and short-duration afterhyperpolarization (46). Most cells also expressed mRNA transcripts for parvalbumin (*Pvalb)* or parathyroid hormone like hormone (*Pthlh*), two markers of striatal FSIs (47, 48) (**Figure S1D,E**).

Given the sparse distribution of FSIs in the NAc, we used fiber photometry to collect bulk calcium signal from FSI ensembles in freely behaving mice (49, 50). To establish the sensitivity of this approach, PV-2A-Cre mice were stereotaxically co-injected with AAVs expressing two Cre-dependent constructs (Figure 2A): the genetically-encoded calcium indicator jGCaMP7s (51), and the red-shifted excitatory opsin ChrimsonR (52). To deliver excitation light and collect emitted fluorescence, a fiber-optic was chronically implanted above the site of virus injection (**Figure S2**). Immunohistochemical examination of infected brain slices revealed sparse and overlapping viral co-expression beneath the fiber-optic implant (Figure 2B). We tethered freely moving mice to the fiber photometry system and delivered amber light (595 nm) through the fiber-optic implant to activate ChrimsonR (Figure 2C). We detected robust jGCaMP7s signals that varied in a dose-dependent fashion with stimulation frequency (F_1.94,9.67_=6.77, p=0.015; Figure 2D), pulse number (F_3.99,19.98_=4.92, p=0.006; Figure 2E), and light power (F_1.89,9.47_=14.44, p=0.001; Figure 2F).

**Figure 2.**
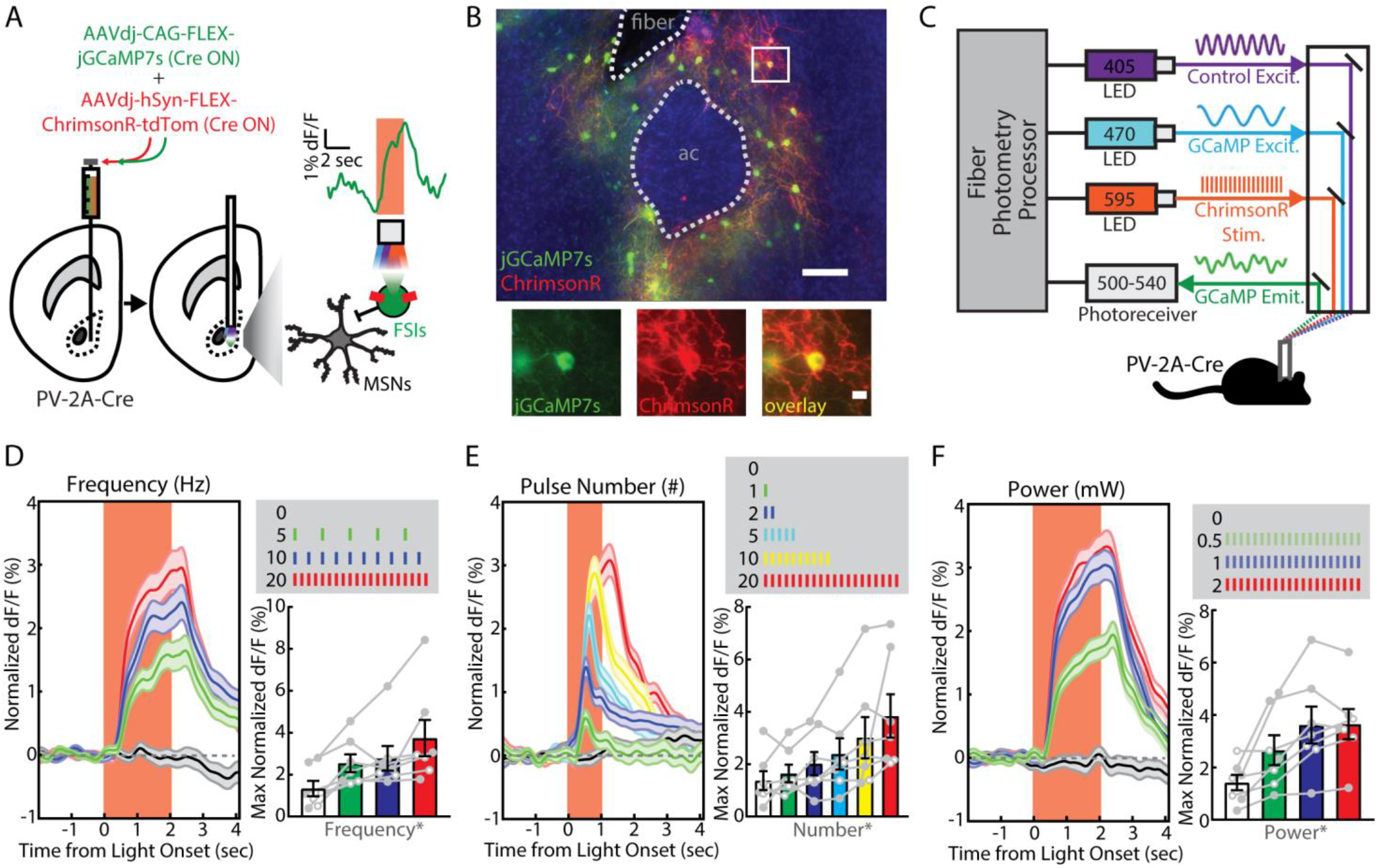
Optogenetically-evoked calcium signals from fast-spiking interneurons (FSIs) in the nucleus accumbens (NAc) core. (**A**) Viral injection of Cre-dependent ChrimsonR and jGCaMP7s into the NAc core of PV-2A-Cre mice (n = 7; 4 male, 3 female). Inset shows photometry signal on a single representative trial. (**B**) (**left**) Visualization of fiber photometry implant placed above FSIs co-expressing (red) ChrimsonR and (green) jGCaMP7s (ac, anterior commissure; scale bar, 100 µm). (**right**) Magnified image of a single FSI (scale bar, 10 µm). (**C**) Setup for simultaneous in vivo optogenetic light stimulation and fiber photometry monitoring. Optogenetically-evoked jGCaMP7s signal measured in vivo by varying (**D**) stimulation frequency (0, 5, 10, 20 Hz; n = 20 trials each), (**E**) pulse number (0, 1, 2, 5, 10, 20; n = 20 trials each), or (**F**) light power (0, 0.5, 1, 2 mW; n = 20 trials each). *p<0.05 for the indicated main effect.

### Fiber Photometry Monitoring of FSIs in the 5-CSRTT

After verifying the specificity and reliability of fiber photometry signals from FSIs in the NAc, we proceeded to record these signals while mice performed the 5-CSRTT. Experimental mice included those with validated jGCaMP7s expression from the preceding optogenetic experiments (see Figure 2), as well as a control cohort of PV-2A-Cre mice expressing Cre-dependent eYFP. We collected fiber photometry data across five consecutive sessions in the final stage of training, with simultaneous excitation of jGCaMP7s at a calcium-dependent wavelength (470 nm), as well as an isosbestic wavelength (405 nm) used to correct for bleaching and movement artifact (Figure 3A). The fiber photometry signal was time-locked to different task events using TTL outputs from the behavioral control software (Figure 3B,C), and normalized to a 2 second baseline period preceding trial onset.

**Figure 3.**
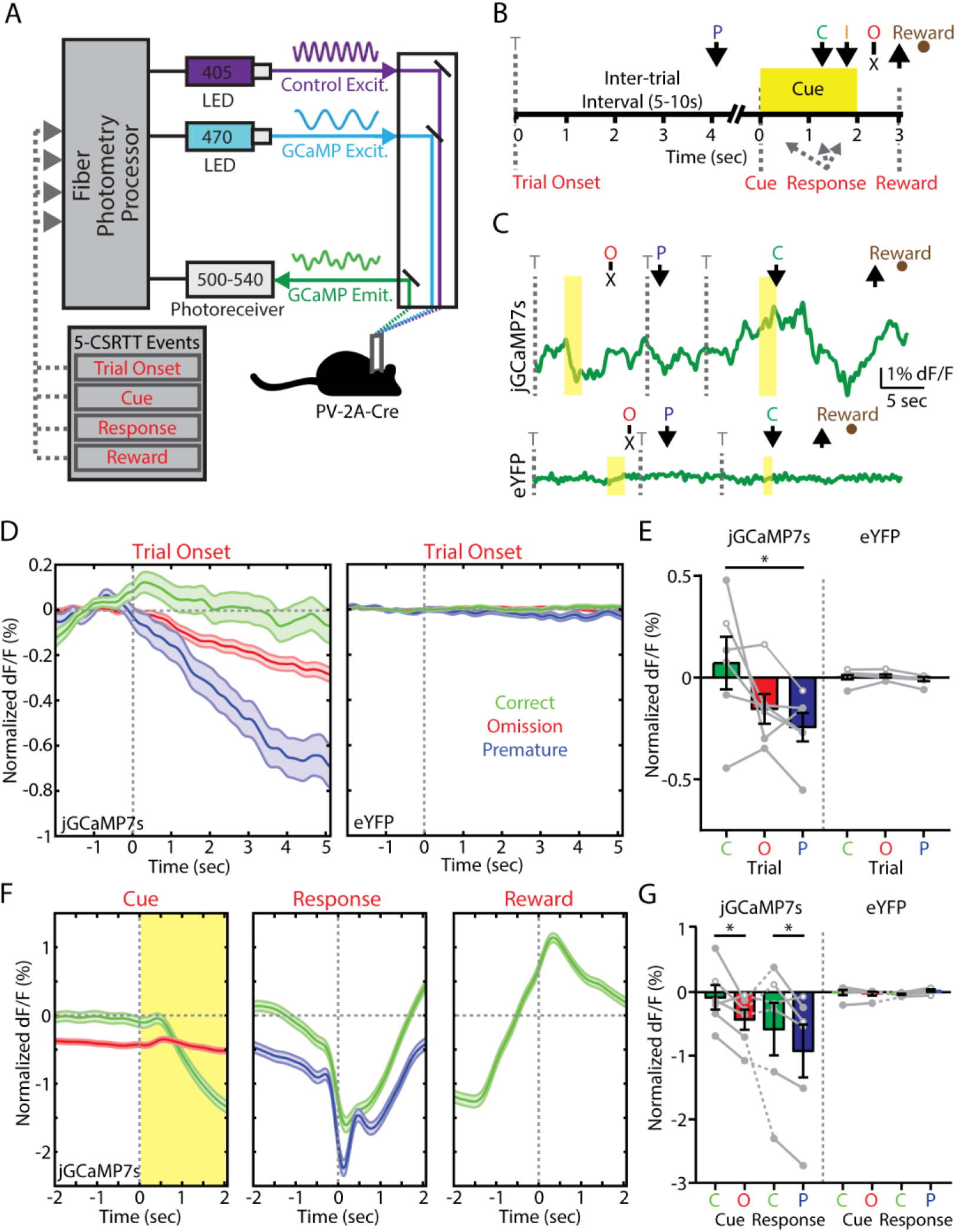
Fast-spiking interneuron (FSI) activity within the nucleus accumbens (NAc) core predicts trial outcome in the 5-choice serial reaction time task (5-CSRTT). (**A**) Fiber photometry setup for in vivo monitoring of jGCaMP7s expressed by FSIs during the 5-CSRTT. (**B**) Timeline of the 5-CSRTT, including events for fiber photometry analysis. Arrows indicate nose poke response. T, trial onset; P, premature; C, correct; I, incorrect; O, omission. (**C**) Example fiber photometry signal across various trial types in PV-2A-Cre mice expressing (**top**) jGCaMP7s or (**bottom**) eYFP. (**D**) Fiber photometry signal aligned to trial onset for mice expressing (**left**) jGCaMP7s (trials: correct, 919; premature, 425; omission, 4871) or (**right**) eYFP (trials: correct, 1336; premature, 305; omission, 3697). (**E**) Fiber photometry data averaged within animal for jGCaMP7s (mice: n = 6: 3 male, 3 female) or eYFP (mice: n = 6: 3 male, 3 female). (**F**) Fiber photometry signal from mice expressing jGCaMP7s aligned to (**left)** cue, (**middle**) nose poke response, or (**right**) magazine entry for food reward. (**G**) Fiber photometry data averaged within animal. Bar graphs represent normalized dF/F values averaged over two seconds following (trial onset) or prior to (cue, response) each aligned event. *p<0.05, LSD post-hoc test.

We first examined fiber photometry signals aligned to trial onset, which coincided with the beginning of the ITI period (Figure 3D, **left**). FSI activity was sustained throughout the ITI on correct trials, whereas it declined on both omission and premature trials. These response profiles were absent in control mice expressing eYFP (Figure 3D, **right**), indicating the jGCaMP7s signal was not a motion artifact. After averaging calcium signals for each animal during the 2 seconds after trial onset, there was a significant Trial x Group interaction (F_2,16_=5.42, p=0.016), with a large effect size in both females 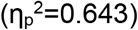 and males 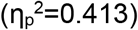. Mice expressing jGCaMP7s showed a significant difference in FSI activity following the onset of correct versus premature trials (p<0.05, LSD post-hoc test; Figure 3E).

Because of the variable interval from trial onset to cue presentation, response, and reward retrieval, we also aligned the fiber photometry signal to these other task events. FSI activity on correct trials was sustained before cue presentation (Figure 3F, **left**) as well as response (Figure 3F, **middle**), and increased when mice retrieved reward (Figure 3F, **right**). The response signal did not vary as a function of spatial location relative to implanted hemisphere (**Figure S3**). After averaging calcium signals for each animal during the 2 seconds preceding each event (Figure 3G), we observed significantly elevated FSI activity before cue presentation on correct versus omission trials (Trial x Group: F_1,8_=10.41, p=0.012), and before the response on correct versus premature trials (Trial x Group: F_1,8_=9.34, p=0.016). Incorrect responses were not analyzed, as they represented <2% of all trials. Altogether, these data show reduced FSI activity predicts poor behavioral performance in the form of premature responses.

### Effects of FSI Manipulations on Projection Neurons

Given the relationship between FSI activity and behavioral performance in the 5-CSRTT, our next goal was to determine the effects of directly manipulating FSI activity *in vivo*. We assessed the efficacy of optogenetic and chemogenetic FSI manipulations by analyzing interactions with medium spiny projection neurons (MSNs) in the NAc core. First, we used red-shifted optogenetic actuators to manipulate FSI activity while recording GCaMP signal from MSNs with fiber photometry (Figure 4A,B). Stimulation of FSIs expressing ChrimsonR decreased the GCaMP signal from MSNs (Figure 4C). To optogenetically inhibit FSIs, we expressed halorhodopsin (eNpHR3.0), a chloride pump that produces robust neuronal hyperpolarization (33, 53, 54). Inhibition of FSIs expressing halorhodopsin increased GCaMP signal from MSNs (Figure 4D), with a robust difference between the effects of optogenetic activation and inhibition of FSIs (F_1,6_=17.37, p=0.006; Figure 4E).

**Figure 4.**
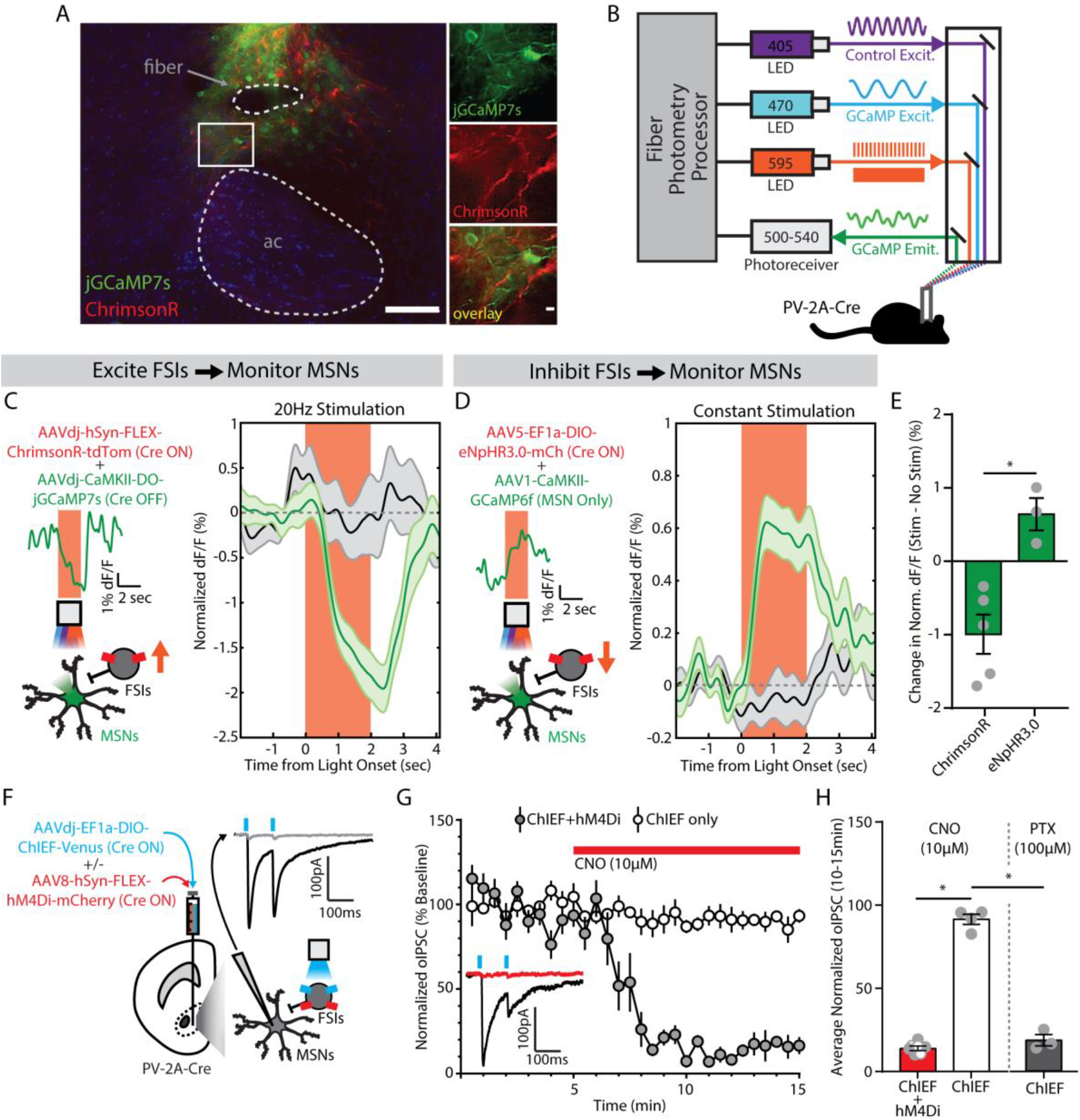
Fast-spiking interneurons (FSIs) inhibit medium spiny neurons (MSNs) in the nucleus accumbens (NAc) core. (**A**) (**left**) Visualization of fiber photometry implant placed above FSIs expressing ChrimsonR (red) and MSNs expressing jGCaMP7s (green; ac, anterior commissure; scale bar, 100 µm). (**right**) Magnified image of FSI-MSN juxtaposition (scale bar, 10 µm). (**B**) Setup for simultaneous red-shifted optogenetic manipulation (pulsed for ChrimsonR, constant for eNpHR3.0) and fiber photometry monitoring in vivo. (**C**) (**left**) Viral co-injection of Cre-dependent ChrimsonR to excite FSIs with Cre-inactivated jGCaMP7s to monitor MSNs. Insets show photometry signal on a single representative trial. (**right**) MSN signal decreases from baseline during FSI excitation (green, n=20 trials/mouse), compared to no stimulation trials (black, n=20 trials/mouse) (n=5 male mice). (**D**) (**left**) Viral co-injection of Cre-dependent NpHR3.0 to inhibit FSIs with CaMKII-promoted GCaMP6f to monitor MSNs. Insets show photometry signal on a single representative trial. (**right**) MSN signal increases from baseline during FSI inhibition (green, n=20 trials/mouse), compared to no stimulation trials (black, n=20 trials/mouse) (n=3 male mice). (**E**) Significant difference in MSN signal between FSI excitation and inhibition. *p<0.05, main effect of group. (**F**) Viral co-injection of Cre-dependent ChIEF, with or without Cre-dependent hM4Di. Inset shows an optogenetically-evoked inhibitory postsynaptic current (oIPSC, black) blocked by picrotoxin (100 µM, gray). (**G)** oIPSCs in slices co-expressing hM4Di were reduced by bath application of clozapine-N-oxide (CNO, 10µM, red bar) (ChIEF only, n=4 male cells; ChIEF+hM4Di, n=7 cells; 3 female, 4 male). Inset shows an example trace before (black) and after (red) CNO, from a slice expressing ChIEF+hM4Di. (**H**) oIPSCs were significantly reduced by chemogenetic inhibition (red) or picrotoxin (PTX, gray; n=3 male cells). *p<0.05, main effect of group.

We also measured synaptic interactions between FSIs and MSNs using whole-cell patch-clamp electrophysiology. To stimulate FSIs and their synaptic outputs, we injected PV-2A-Cre mice with a Cre-dependent AAV expressing ChIEF, an engineered hybrid of channelrhodopsin-1 and channelrhodopsin-2 with optimized biophysical properties (55, 56). We then prepared acute brain slices and performed voltage-clamp recordings from uninfected MSNs, evoking optical inhibitory post-synaptic currents (oIPSCs) with blue light pulses (Figure 4F). Some PV-2A-Cre mice were co-injected with Cre-dependent AAVs expressing ChIEF as well as a Gi-coupled chemogenetic actuator, hM4Di (57). In these mice, activation of hM4Di with clozapine-N-oxide (CNO) substantially reduced oIPSC amplitude (F_1,8_=413.73, p<0.001; Figure 4G). In slices from control animals that were not injected with hM4Di, CNO had no effect but oIPSCs were blocked by picrotoxin (100 µM), a GABA_A_ receptor antagonist (F_1,8_=200.10, p<0.001; Figure 4H).

### Chemogenetic Inhibition of FSIs Increases Impulsivity

To translate this chemogenetic manipulation of FSI activity to behavioral performance of the 5-CSRTT, PV-2A-Cre mice and wild-type littermates received bilateral injections of an AAV expressing Cre-dependent hM4Di (Figure 5A), with infection largely confined to the NAc core (**Figure S4**). Current-clamp recordings confirmed that CNO (10 µM) decreased the intrinsic excitability of infected FSIs in acute brain slices (Figure 5B), significantly reducing maximum firing rate and hyperpolarizing the resting membrane potential (**Figure S5**). When mice reached the last stage of training, we injected saline or CNO (2 mg/kg) 30 minutes before testing on separate days, followed by tests of impulsivity and attention (see Figure 1) under the same conditions on subsequent days.

**Figure 5.**
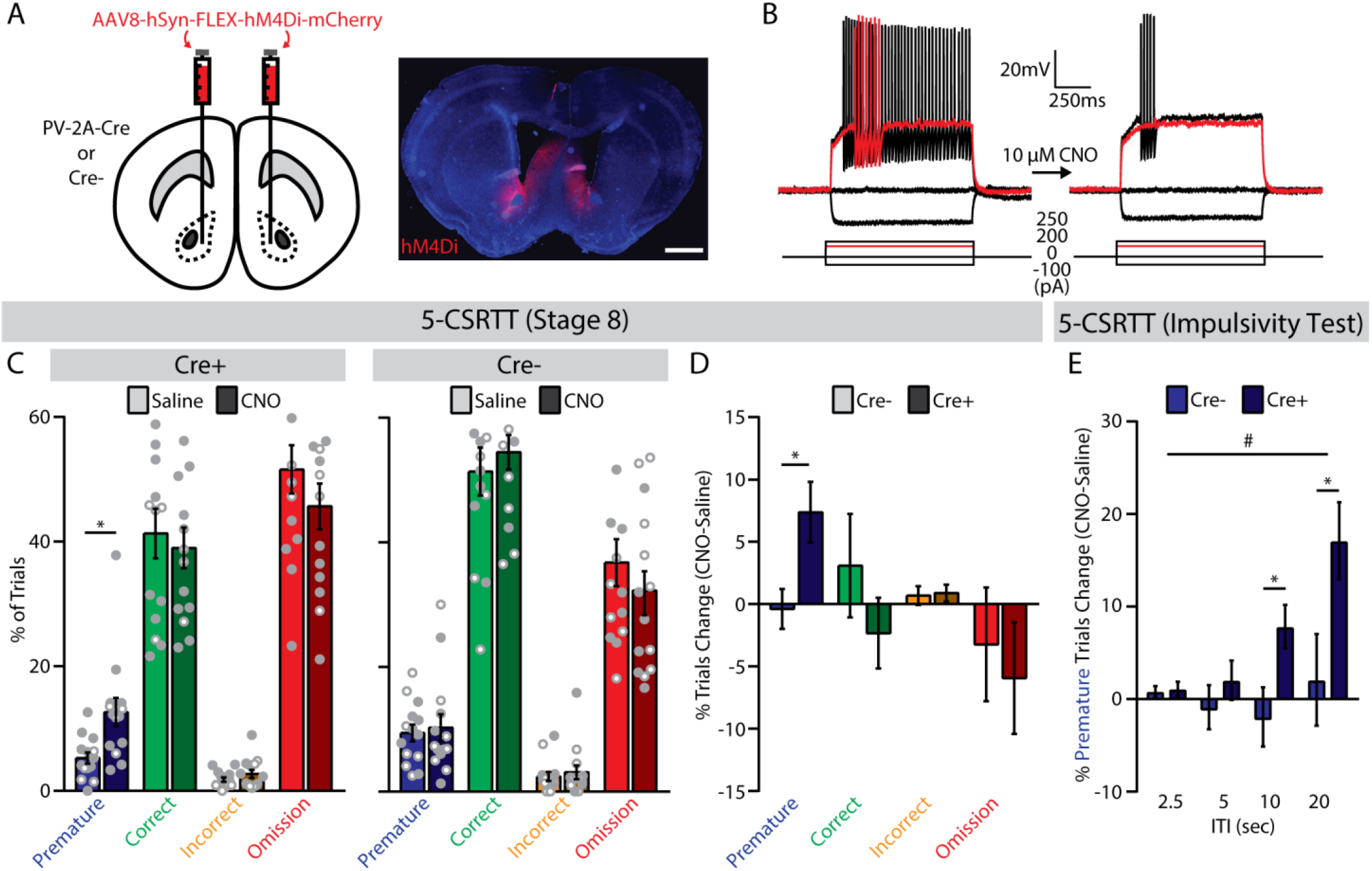
Chemogenetic inhibition of fast-spiking interneurons (FSIs) increases premature responses in the 5-choice serial reaction time task (5-CSRTT). (**A**) (**left**) Bilateral viral injection of Cre-dependent hM4Di into the NAc core of PV-2A-Cre or wild-type (Cre-) mice, and (**right**) histological verification of hM4Di expression (scale bar, 1 mm). (**B**) Whole-cell current-clamp recording of a representative FSI before (**left**) and after (**right**) bath application of CNO (10 µM). Red trace indicates lowest input current needed to elicit spiking (rheobase) before CNO application. (**C**) Trial outcomes for the final stage of the 5-CSRTT in (**left**) PV-2A-Cre (n=14, 10 male, 4 female) and (**right**) wild-type (n=14; 7 male, 7 female) mice following injection of saline or CNO (2 mg/kg). *p<0.05, main effect of treatment. (**D**) Systemic administration of CNO (2 mg/kg) selectively increased the percentage of premature trials in PV-2A-Cre mice expressing hM4Di. *p<0.05, main effect of group. (**E**) In the impulsivity test, systemic administration of CNO (2 mg/kg) increased the percentage of premature trials in PV-2A-Cre mice, especially at longer ITIs. *p<0.05, LSD post-hoc test. #p<0.05, group x ITI interaction.

In PV-2A-Cre mice, there was no change in total trial number after injection of CNO versus saline (**Table S2**). However, the percentage of premature trials increased significantly after injection of CNO versus saline (F_1,12_=7.15, p=0.020; Figure 5C, **left; Figure S6A**), with large effect sizes in both females 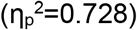 and males 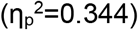. This effect was not observed in wild-type littermates (F_1,12_ <1; Figure 5C, **right; Figure S6A**), demonstrating dependence on hM4Di expression. Comparison of difference scores (CNO-saline) between genotypes confirmed a specific effect on premature responses (F_1,24_=6.39, p=0.018), with no significant change in other trial types (Figure 5D) or the number of perseverative responses (PV-2A-Cre: 1.71±1.76; wild-type: 2.86±1.13; F_1,24_<1). Importantly, chemogenetic inhibition of FSIs did not affect latencies to respond or retrieve reward (**Figure S6B,C**), nor did it alter locomotor activity in an open field (**Figure S7**)

In the impulsivity test (**Figure S8**), CNO administration increased premature responses in PV-2A-Cre mice when the ITI was relatively long. This pattern was evident in difference scores, which revealed a significant Group x ITI interaction (F_1,24_ =3.82, p=0.043), with no effect of CNO in the wild-type control group (Figure 5E). In the attention test (**Figure S9**), there was an increase in correct trials following CNO injection in PV-2A-Cre mice, but no significant changes in other trial types or differences with wild-type littermates.

### Temporally Delimited Optogenetic Inhibition of FSIs Increases Impulsivity

The preceding chemogenetic manipulation was effective, but lacks the temporal precision of our fiber photometry recordings, which showed sustained FSI activity on correct trials during the ITI. To provide temporally precise inhibition of FSI activity, we used a halorhodopsin-based optogenetic strategy (33) already validated with fiber photometry (see Figure 4D). PV-2A-Cre mice received bilateral injections of an AAV expressing Cre-dependent eNpHR3.0 or eYFP (Figure 6A), followed by chronic implantation of fiber-optic cannulae above the sites of virus injection in the NAc core (**Figure S10**). Current-clamp recordings in acute brain slices confirmed that amber light (593 nm) hyperpolarized infected FSIs, effectively preventing and interrupting action potentials (Figure 6B,C).

**Figure 6.**
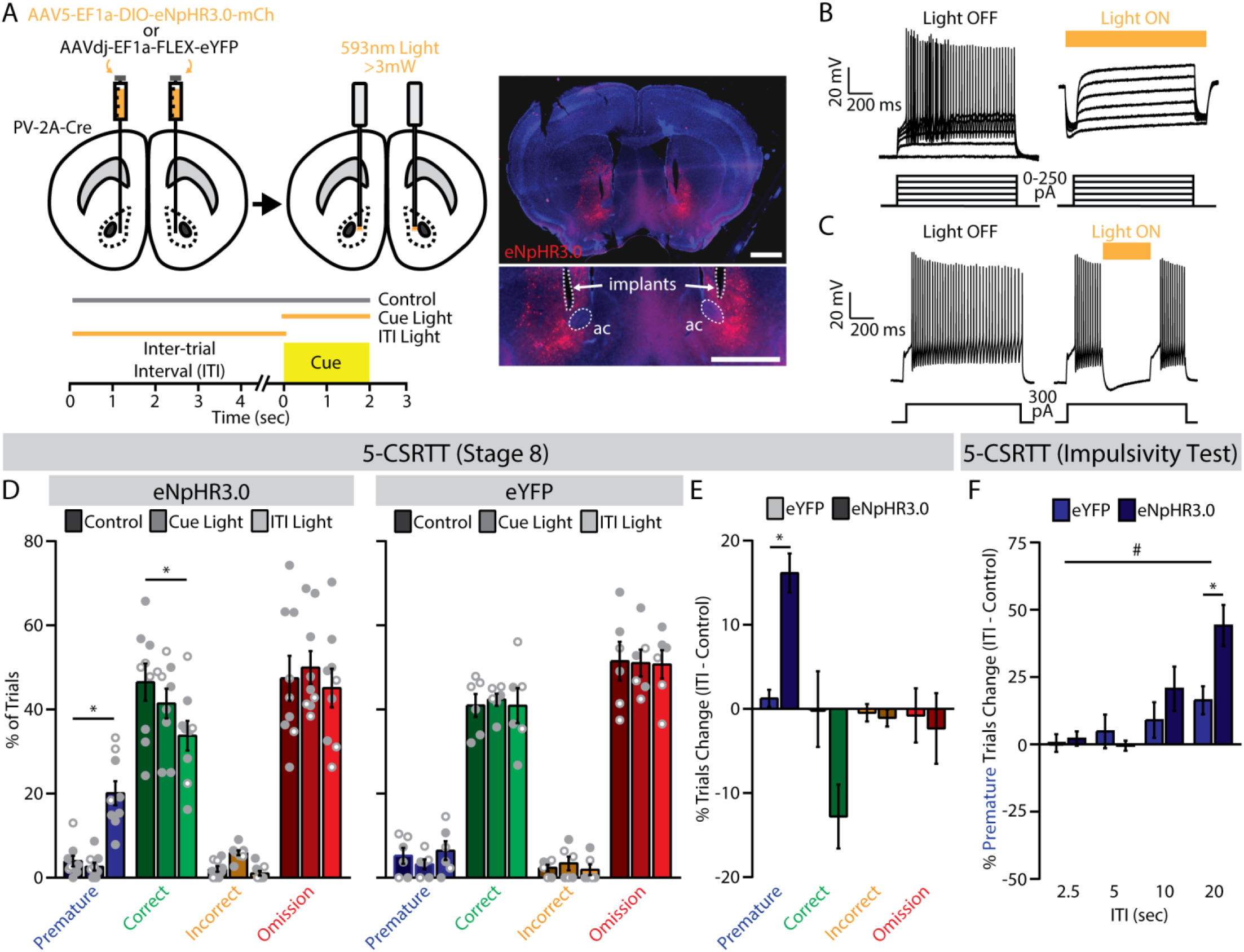
Optogenetic silencing of fast-spiking interneurons (FSIs) during the inter-trial interval (ITI) increases impulsive behavior in the 5-choice serial reaction time task (5-CSRTT). (**A**) (**top**) Bilateral viral injection of Cre-dependent eNpHR3.0 (n=9; 6 male, 3 female) or eYFP (n=6; 3 male, 3 female) into the NAc core of PV-2A-Cre mice; (**bottom**) trial types for 5-CSRTT testing; (**right**) histological verification of fiber-optic placement and eNpHR3.0 expression (ac, anterior commissure; scale bars, 1 mm). (**B**) Whole-cell current-clamp recording of a representative FSI expressing NpHR3.0 (**left**) before and (**right**) after optogenetic inhibition. (**C**) Light interrupts spiking of a FSI. (**D**) Trial outcomes for the final stage of the 5-CSRTT in PV-2A-Cre mice expressing (**left**) eNpHR3.0 (n=9; 6 male, 3 female) or (**right**) eYFP (n=6; 3 male, 3 female) across the three trial types. *p<0.05, LSD post-hoc test. (**E**) Optogenetic inhibition during the ITI significantly increased the percentage of premature trials in eNpHR3.0-expressing mice. *p<0.05, main effect of group. (**F**) During the impulsivity test, optogenetic inhibition during the ITI increased the percentage of premature trials in eNpHR3.0-expressing mice, especially at longer ITIs. *p<0.05, LSD post-hoc test. #p<0.05, group x ITI interaction.

When mice reached the last stage of training, we conducted test sessions with interleaved presentation of three trial conditions: light delivery throughout the ITI, light delivery during cue presentation, or no light (i.e., control) (Figure 6A, **bottom**). In mice expressing eNpHR3.0 in NAc FSIs (Figure 6D, **left)**, light delivery throughout the ITI significantly increased the percentage of premature trials (F_2.14_=65.60, p<0.001) and decreased the percentage of correct trials (F_2,14_=5.11, p=0.022), relative to the dark control condition (p<0.05, LSD post-hoc test). Light delivery during the cue had no effect in mice expressing eNpHR3.0, and the eYFP control group showed no significant effect of light delivery under any condition (Figure 6D; **Figure S11A,B**). Total trial number was similar in eNpHR3.0 and eYFP groups (**Table S3**). Comparison of difference scores (ITI-control) between groups confirmed a specific effect on premature responses in mice expressing eNpHR3.0 (F_1,11_=34.86, p<0.001), with large effect size in both females 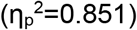 and males 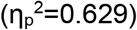, and no significant change in other trials types (Figure 6E). Optogenetic inhibition of FSIs also did not change latencies to respond or retrieve reward (**Figure S11C,D**).

To corroborate the specificity of these effects, we performed optogenetic manipulation in the same cohort of mice during tests of impulsivity and attention. We adopted a simplified trial structure that involved randomized light delivery during the ITI for the impulsivity test or during the cue for the attention test, and compared either trial type to interleaved control trials. In the attention test, there were significant Group x Cue interactions for correct trials (F_3,33_=3.62, p=0.023) and omissions (F_3,33_=4.55, p=0.009), with different effects at 2.5- and 5-second cue durations (**Figure S12**). A clearer pattern emerged in the impulsivity test, where a significant Group × ITI interaction (F_3,33_=3.14, p=0.038) indicated that optogenetic inhibition of FSIs caused more premature responses at longer ITIs (Figure 6F), as well as fewer correct responses (**Figure S13**). Collectively, these data indicate sustained activity of FSIs in the NAc core is necessary to constrain premature responses in the 5-CSRTT, and that disruption of FSI function can lead to impulsive behavior.

## DISCUSSION

Decades of research clearly implicate the NAc in controlling multiple facets of impulsivity, but the contribution of specific cell types within this heterogeneous brain region has remained unclear. In this study, we focused on the contribution of FSIs, which are few in number but exert powerful inhibitory control over MSN output from the NAc (22–25). We used PV-2A-Cre transgenic mice to monitor and manipulate the activity of FSIs in the NAc core. Our data suggest the sustained activity of these cells positively correlates with successful control of impulsive behavior in the 5-CSRTT. Furthermore, using chemogenetic and optogenetic methods to inhibit FSIs in the NAc core, we observed more impulsive behavior across a range of task conditions. These findings indicate FSIs in the NAc core play a key role in impulse control.

The sparse distribution of FSIs in the NAc has historically made these cells difficult to study. FSIs represent fewer than 1% of all neurons in the dorsal striatum of rodents (58), with immunohistochemical and electrophysiological evidence suggesting FSIs are even less abundant in the NAc (59, 60). Several aspects of our experimental design facilitated analysis of these scarce interneurons, including bulk calcium imaging of NAc tissue with fiber photometry and jGCaMP7s, a recently developed and highly sensitive genetically encoded calcium indicator (51). This approach enabled real-time monitoring of FSI activity in the NAc core as mice performed the 5-CSRTT. On correct trials, FSI activity was sustained during the ITI, then declined after cue presentation and nose poke in the appropriate location. In contrast, FSI activity declined during the ITI on premature trials, and remained low at the time of premature response. To move beyond correlations between FSI activity and behavior, we used chemogenetic and optogenetic approaches to specifically inhibit FSIs in the NAc core. Both manipulations increased the frequency of premature responses, directly linking FSI activity with successful impulse control. Importantly, CNO injection and light delivery had no effect in control animals, demonstrating specificity in the behavioral effects of these manipulations.

Compared to these effects on impulsivity, manipulations of NAc FSIs had no consistent effect on sustained attention in the 5-CSRTT. Chemogenetic inhibition tended to reduce omissions in the impulsivity and attention tests, with a corresponding increase in correct responses. However, these trends were not apparent under standard task conditions (Stage 8) or with optogenetic inhibition. The chronic nature of chemogenetic FSI inhibition could provoke compensatory changes in GABA release from other sources, such as low-threshold spiking interneurons or local MSN collaterals (61). The attention and impulsivity tests introduce novel task conditions, which may enhance recruitment of low-threshold spiking interneurons (62), leading to co-release of GABA and other neuromodulators (nitric oxide, somatostatin, neuropeptide Y) that affect attention. Regardless of the explanation for this inconsistency, our findings contrast with studies of FSIs in prefrontal cortex, which are also active during the ITI but appear to sustain attention in the 5-CSRTT (32).

Our data show FSIs form strong inhibitory synaptic connections onto MSNs in the NAc, as previously reported in brain slices (22–25). By combining optogenetics and fiber photometry, we found that FSIs in the NAc core negatively modulate calcium signals from surrounding MSNs, consistent with a net inhibitory influence of NAc FSIs on MSN activity *in vivo*. The increased impulsivity observed after chemogenetic or optogenetic FSI inhibition is thus likely to involve disinhibition of NAc output from MSNs. We speculate that the amount of GABA released by FSIs may be a key factor controlling impulsivity, since highly impulsive rats have decreased NAc GABA levels (19), but no apparent change in the number of GABA receptors (63). Local MSN collaterals are another potential source of GABA release (61), but highly impulsive rats have lower expression of glutamate decarboxylase in the NAc core (18), and this key GABA synthetic enzyme is encoded by two genes (*Gad1* and *Gad2*) highly expressed by FSIs (64). Reduced GABA synthesis by FSIs may thus contribute to increased impulsivity observed after local knockdown of glutamate decarboxylase expression in the NAc core (18).

Our data are consistent with evidence that FSI activity in the dorsal striatum can delay goal-directed actions (65). Other evidence suggests these cells suppress inappropriate movements and actions: striatal FSIs are lost in Tourette’s syndrome (66), and experimental disruptions of FSIs in the rodent striatum produce abnormal behaviors, including motor stereotypies and dyskinesias (67–71). These multiple functions are consistent with the fact that FSIs exert a strong inhibitory influence over both striatonigral and striatopallidal MSNs (28, 72), pointing to a multifaceted role in regulating action selection and choice execution (73, 74). Regulation of learning (33, 34) and habit formation (35) likely involves plasticity of inhibitory FSI synapses onto MSNs, as well as plasticity of excitatory synaptic inputs to FSIs themselves (23, 75). However, maladaptive engagement of FSI plasticity could contribute to addiction and other disease states (36, 37, 76). The present findings thus have broad implications for understanding addiction and other neuropsychiatric disorders associated with impulsivity and associated dysfunction of limbic neurocircuitry (2, 8, 11, 77).

## ACKNOWLEDGEMENTS

Research reported in this publication was supported by the University of Minnesota’s MnDRIVE (Minnesota’s Discovery, Research and Innovation Economy) initiative (to MTP, BHT, and PER); grants from the National Institutes of Health (F32MH118794 to MTP and R00DA037279 to PER); and the Whitehall Foundation, a NARSAD Young Investigator Grant from the Brain & Behavior Research Foundation, and an MQ Fellows Award (all to PER). The University of Minnesota MnDRIVE Optogenetics Core provided technical support for fiber photometry experiments. Some of the viral vectors used in this study were generated by the University of Minnesota Viral Vector and Cloning Core, as well as the Stanford University Gene Virus and Vector Core. We thank Justin Lines for fiber photometry analysis troubleshooting; Eshaan Iyer for early piloting of the 5-CSRTT; Sowmya Narayan and Sumyuktha Vijay for assistance with behavioral experiments; Drs. Rocio Gomez-Pastor and Esther Krook-Magnuson for sharing advice and reagents; and Carlee Toddes and Dieter Brandner for helpful discussion of the manuscript. A preprint version of this manuscript is available on bioRxiv: https://doi.org/10.1101/516609.

The authors report no biomedical financial interests or potential conflicts of interest.

## REFERENCES

1. Dalley JW, Everitt BJ, Robbins TW (2011): Impulsivity, compulsivity, and top-down cognitive control. Neuron. 69: 680–694.

2. Dalley JW, Robbins TW (2017): Fractionating impulsivity: Neuropsychiatric implications. Nat Rev Neurosci. 18: 158–171.

3. Verdejo-García A, Lawrence AJ, Clark L (2008): Impulsivity as a vulnerability marker for substance-use disorders: Review of findings from high-risk research, problem gamblers and genetic association studies. Neurosci Biobehav Rev. 32: 777–810.

4. Winstanley CA, Eagle DM, Robbins TW (2006): Behavioral models of impulsivity in relation to ADHD: Translation between clinical and preclinical studies. Clin Psychol Rev. 26: 379–395.

5. Chamberlain SR, Robbins TW, Winder-Rhodes S, Mller U, Sahakian BJ, Blackwell AD, Barnett JH (2011): Translational approaches to frontostriatal dysfunction in attention-deficit/hyperactivity disorder using a computerized neuropsychological battery. Biol Psychiatry. 69: 1192–1203.

6. Javdani S, Sadeh N, Verona E (2011): Suicidality as a function of impulsivity, callous-unemotional traits, and depressive symptoms in youth. J Abnorm Psychol. 120: 400–413.

7. Basar K, Sesia T, Groenewegen H, Steinbusch HWM, Visser-Vandewalle V, Temel Y (2010): Nucleus accumbens and impulsivity. Prog Neurobiol. 92: 533–557.

8. Morris LS, Kundu P, Baek K, Irvine MA, Mechelmans DJ, Wood J, et al. (2016): Jumping the gun: Mapping neural correlates of waiting impulsivity and relevance across alcohol misuse. Biol Psychiatry. 79: 499–507.

9. Kable JW, Glimcher PW (2007): The neural correlates of subjective value during intertemporal choice. Nat Neurosci. 10: 1625–1633.

10. McClure SM (2004): Separate neural systems value immediate and delayed monetary rewards. Science (80-). 306: 503–508.

11. Carmona S, Proal E, Hoekzema EA, Gispert JD, Picado M, Moreno I, et al. (2009): Ventro-striatal reductions underpin symptoms of hyperactivity and impulsivity in attention-deficit/hyperactivity disorder. Biol Psychiatry. 66: 972–977.

12. Cardinal RN, Pennicott DR, Lakmali C, Robbins TW, Everitt BJ (2001): Impulsive choice induced in rats by lesions of the nucleus accumbens core. Science (80-). 292: 2499–2501.

13. Christakou A (2004): Prefrontal cortical-ventral striatal interactions involved in affective modulation of attentional performance: implications for corticostriatal circuit function. J Neurosci. 24: 773–780.

14. Pothuizen HHJ, Jongen-Rêlo AL, Feldon J, Yee BK (2005): Double dissociation of the effects of selective nucleus accumbens core and shell lesions on impulsive-choice behaviour and salience learning in rats. Eur J Neurosci. 22: 2605–2616.

15. Murphy ER, Robinson ESJ, Theobald DEH, Dalley JW, Robbins TW (2008): Contrasting effects of selective lesions of nucleus accumbens core or shell on inhibitory control and amphetamine-induced impulsive behaviour. Eur J Neurosci. 28: 353–363.

16. Koskinen T, Sirviö J (2001): Studies on the involvement of the dopaminergic system in the 5-HT2agonist (DOI)-induced premature responding in a five-choice serial reaction time task. Brain Res Bull. 54: 65–75.

17. Pezze MA, Dalley JW, Robbins TW (2007): Differential roles of dopamine D1 and D2 receptors in the nucleus accumbens in attentional performance on the five-choice serial reaction time task. Neuropsychopharmacology. 32: 273–283.

18. Caprioli D, Sawiak SJ, Merlo E, Theobald DEH, Spoelder M, Jupp B, et al. (2014): Gamma aminobutyric acidergic and neuronal structural markers in the nucleus accumbens core underlie trait-like impulsive behavior. Biol Psychiatry. 75: 115–123.

19. Sawiak SJ, Jupp B, Taylor T, Caprioli D, Carpenter TA, Dalley JW (2016): In vivo γ-aminobutyric acid measurement in rats with spectral editing at 4.7T. J Magn Reson Imaging. 43: 1308–1312.

20. Hayes DJ, Jupp B, Sawiak SJ, Merlo E, Caprioli D, Dalley JW (2014): Brain γ-aminobutyric acid: A neglected role in impulsivity. Eur J Neurosci. 39: 1921–1932.

21. Hu H, Gan J, Jonas P (2014): Fast-spiking, parvalbumin+ GABAergic interneurons: From cellular design to microcircuit function. Science (80-). 345. doi: 10.1126/science.1255263.

22. Scudder SL, Baimel C, Macdonald EE, Carter AG (2018): Hippocampal-evoked feed-forward inhibition in the nucleus accumbens. J Neurosci. 38: 1971–18.

23. Wright WJ, Schlüter OM, Dong Y (2017): A feedforward inhibitory circuit mediated by CB1-expressing fast-spiking interneurons in the nucleus accumbens. Neuropsychopharmacology. 42: 1146–1156.

24. Taverna S, Canciani B, Pennartz CMA (2007): Membrane properties and synaptic connectivity of fast-spiking interneurons in rat ventral striatum. Brain Res. 1152: 49–56.

25. Winters BD, Kruger JM, Huang X, Gallaher ZR, Ishikawa M, Czaja K, et al. (2012): Cannabinoid receptor 1-expressing neurons in the nucleus accumbens. Proc Natl Acad Sci. 109: E2717–E2725.

26. Koós T, Tepper JM (1999): Inhibitory control of neostriatal projection neurons by GABAergic interneurons. Nat Neurosci. 2: 467–472.

27. Straub C, Saulnier JL, Bègue A, Feng DD, Huang KW, Sabatini BL (2016): Principles of synaptic organization of GABAergic interneurons in the striatum. Neuron. 92: 84–92.

28. Gittis AH, Nelson AB, Thwin MT, Palop JJ, Kreitzer AC (2010): Distinct roles of GABAergic interneurons in the regulation of striatal output pathways. J Neurosci. 30: 2223–34.

29. Cardin JA, Carlén M, Meletis K, Knoblich U, Zhang F, Deisseroth K, et al. (2009): Driving fast-spiking cells induces gamma rhythm and controls sensory responses. Nature. 459: 663–667.

30. Royer S, Zemelman B V., Losonczy A, Kim J, Chance F, Magee JC, Buzsáki G (2012): Control of timing, rate and bursts of hippocampal place cells by dendritic and somatic inhibition. Nat Neurosci. 15: 769–775.

31. Sohal VS, Zhang F, Yizhar O, Deisseroth K (2009): Parvalbumin neurons and gamma rhythms enhance cortical circuit performance. Nature. 459: 698–702.

32. Kim H, Ährlund-Richter S, Wang X, Deisseroth K, Carlén M (2016): Prefrontal parvalbumin neurons in control of attention. Cell. 164: 208–218.

33. Owen SF, Berke JD, Kreitzer AC (2018): Fast-spiking interneurons supply feedforward control of bursting, calcium, and plasticity for efficient learning. Cell. 172: 683–695.e15.

34. Lee K, Holley SM, Shobe JL, Chong NC, Cepeda C, Levine MS, Masmanidis SC (2017): Parvalbumin interneurons modulate striatal output and enhance performance during associative learning. Neuron. 93: 1451–1463.e4.

35. O’Hare JK, Li H, Kim N, Gaidis E, Ade K, Beck J, et al. (2017): Striatal fast-spiking interneurons selectively modulate circuit output and are required for habitual behavior. Elife. 6: 1–26.

36. Wang X, Gallegos DA, Pogorelov VM, O’Hare JK, Calakos N, Wetsel WC, West AE (2018): Parvalbumin interneurons of the mouse nucleus accumbens are required for amphetamine-induced locomotor sensitization and conditioned place preference. Neuropsychopharmacology. 43: 953–963.

37. Yu J, Yan Y, Li K-L, Wang Y, Huang YH, Urban NN, et al. (2017): Nucleus accumbens feedforward inhibition circuit promotes cocaine self-administration. Proc Natl Acad Sci. 201707822.

38. Robbins TW (2002): The 5-choice serial reaction time task: Behavioural pharmacology and functional neurochemistry. Psychopharmacology (Berl). 163: 362–380.

39. Madisen L, Zwingman TA, Sunkin SM, Oh SW, Zariwala HA, Gu H, et al. (2010): A robust and high-throughput Cre reporting and characterization system for the whole mouse brain. Nat Neurosci. 13: 133–140.

40. Rothwell PE, Hayton SJ, Sun GL, Fuccillo M V., Lim BK, Malenka RC (2015): Input- and output-specific regulation of serial order performance by corticostriatal circuits. Neuron. 88: 345–356.

41. Bari A, Dalley JW, Robbins TW (2008): The application of the 5-choice serial reaction time task for the assessment of visual attentional processes and impulse control in rats. Nat Protoc. 3: 759–767.

42. Grissom NM, Herdt CT, Desilets J, Lidsky-Everson J, Reyes TM (2015): Dissociable deficits of executive function caused by gestational adversity are linked to specific transcriptional changes in the prefrontal cortex. Neuropsychopharmacology. 40: 1353–1363.

43. Krueger DD, Osterweil EK, Chen SP, Tye LD, Mark F, Huganir RL, et al. (2011): Cognitive dysfunction and prefrontal synaptic abnormalities in a mouse model of fragile X syndrome. Proc Natl Acad Sci. 108: 2587–2592.

44. Lerner TN, Shilyansky C, Davidson TJ, Evans KE, Beier KT, Zalocusky KA, et al. (2015): Intact-brain analyses reveal distinct information carried by SNc dopamine subcircuits. Cell. 162: 635–647.

45. Owen SF, Liu MH, Kreitzer AC (2019): Thermal constraints on in vivo optogenetic manipulations. Nat Neurosci. 22: 1061–1065.

46. Kawaguchi Y (1993): Physiological, morphological, and histochemical characterization of three classes of interneurons in rat neostriatum. J Neurosci. 13: 4908–23.

47. Saunders A, Macosko E, Wysoker A, Goldman M, Krienen F, Bien E, et al. (2018): A single-cell atlas of cell types, states, and other transcriptional patterns from nine regions of the adult mouse brain. bioRxiv. doi: 10.1101/299081.

48. Muñoz-Manchado AB, Bengtsson Gonzales C, Zeisel A, Munguba H, Bekkouche B, Skene NG, et al. (2018): Diversity of interneurons in the dorsal striatum revealed by single-cell RNA sequencing and PatchSeq. Cell Rep. 24: 2179–2190.e7.

49. Cui G, Jun SB, Jin X, Pham MD, Vogel SS, Lovinger DM, Costa RM (2013): Concurrent activation of striatal direct and indirect pathways during action initiation. Nature. 494: 238–242.

50. Gunaydin LA, Grosenick L, Finkelstein JC, Kauvar I V., Fenno LE, Adhikari A, et al. (2014): Natural neural projection dynamics underlying social behavior. Cell. 157: 1535–1551.

51. Dana H, Sun Y, Mohar B, Hulse B, Hasseman JP, Tsegaye G, et al. (2019): High-performance calcium sensors for imaging activity in neuronal populations and microcompartments. Nat Methods. 16: 649–657.

52. Klapoetke NC, Murata Y, Kim SS, Pulver SR, Birdsey-Benson A, Cho YK, et al. (2014): Independent optical excitation of distinct neural populations. Nat Methods. 11: 338–346.

53. Zhang F, Wang LP, Brauner M, Liewald JF, Kay K, Watzke N, et al. (2007): Multimodal fast optical interrogation of neural circuitry. Nature. 446: 633–639.

54. Gradinaru V, Zhang F, Ramakrishnan C, Mattis J, Prakash R, Diester I, et al. (2010): Molecular and cellular approaches for diversifying and extending optogenetics. Cell. 141: 154–165.

55. Lin JY, Lin MZ, Steinbach P, Tsien RY (2009): Characterization of engineered channelrhodopsin variants with improved properties and kinetics. Biophys J. 96: 1803–1814.

56. Choi K, Holly E, Davatolhagh MF, Beier KT, Fuccillo M V. (2018): Integrated anatomical and physiological mapping of striatal afferent projections. Eur J Neurosci. 1–14.

57. Armbruster BN, Li X, Pausch MH, Herlitze S, Roth BL (2007): Evolving the lock to fit the key to create a family of G protein-coupled receptors potently activated by an inert ligand. Proc Natl Acad Sci. 104: 5163–5168.

58. Luk KC, Sadikot AF (2001): GABA promotes survival but not proliferation of parvalbumin-immunoreactive interneurons in rodent neostriatum: An in vivo study with stereology. Neuroscience. 104: 93–103.

59. Kita H, Kosaka T, Heizmann CW (1990): Parvalbumin-immunoreactive neurons in the rat neostriatum: a light and electron microscopic study. Brain Res. 536: 1–15.

60. Berke JD, Okatan M, Skurski J, Eichenbaum H (2004): Oscillatory entrainment of striatal neurons in freely moving rats. Neuron. 43: 883–896.

61. Burke DA, Rotstein HG, Alvarez VA (2017): Striatal local circuitry: A new framework for lateral inhibition. Neuron. 96: 267–284.

62. Holly EN, Davatolhagh MF, Choi K, Alabi OO, Vargas Cifuentes L, Fuccillo M V. (2019): Striatal low-threshold spiking interneurons regulate goal-directed learning. Neuron. 103: 1–10.

63. Jupp B, Caprioli D, Saigal N, Reverte I, Shrestha S, Cumming P, et al. (2013): Dopaminergic and GABA-ergic markers of impulsivity in rats: Evidence for anatomical localisation in ventral striatum and prefrontal cortex. Eur J Neurosci. 37: 1519–1528.

64. Saunders A, Macosko EZ, Wysoker A, Goldman M, Krienen FM, de Rivera H, et al. (2018): Molecular diversity and specializations among the cells of the adult mouse brain. Cell. 174: 1015–1030.e16.

65. Tecuapetla F, Jin X, Lima SQ, Costa RM (2016): Complementary contributions of striatal projection pathways to action initiation and execution. Cell. 166: 703–715.

66. Kalanithi PS a, Zheng W, Kataoka Y, DiFiglia M, Grantz H, Saper CB, et al. (2005): Altered parvalbumin-positive neuron distribution in basal ganglia of individuals with Tourette syndrome. Proc Natl Acad Sci U S A. 102: 13307–12.

67. Burguière E, Monteiro P, Feng G, Graybiel AM (2013): Optogenetic stimulation of lateral orbitofronto-striatal pathway suppresses compulsive behaviors. Science (80-). 340: 1243–1246.

68. Gernert M, Hamann M, Bennay M, Löscher W, Richter A, Loscher W, Richter A (2000): Deficit of striatal parvalbumin-reactive GABAergic interneurons and decreased basal ganglia output in a genetic rodent model of idiopathic paroxysmal dystonia. J Neurosci. 20: 7052–7058.

69. Rapanelli M, Frick LR, Xu M, Groman SM, Jindachomthong K, Tamamaki N, et al. (2017): Targeted interneuron depletion in the dorsal striatum produces autism-like behavioral abnormalities in male but not female mice. Biol Psychiatry. 82: 194–203.

70. Xu M, Li L, Pittenger C (2016): Ablation of fast-spiking interneurons in the dorsal striatum, recapitulating abnormalities seen post-mortem in Tourette syndrome, produces anxiety and elevated grooming. Neuroscience. 324: 321–329.

71. Gittis AH, Leventhal DK, Fensterheim BA, Pettibone JR, Berke JD, Kreitzer AC (2011): Selective inhibition of striatal fast-spiking interneurons causes dyskinesias. J Neurosci. 31: 15727–15731.

72. Planert H, Szydlowski SN, Hjorth JJJ, Grillner S, Silberberg G (2010): Dynamics of synaptic transmission between fast-spiking interneurons and striatal projection neurons of the direct and indirect pathways. J Neurosci. 30: 3499–3507.

73. Gage GJ, Stoetzner CR, Wiltschko AB, Berke JD (2010): Selective activation of striatal fast-spiking interneurons during choice execution. Neuron. 67: 466–479.

74. Berke JD (2008): Uncoordinated firing rate changes of striatal fast-spiking interneurons during behavioral task performance. J Neurosci. 28: 10075–10080.

75. Mathur BN, Tanahira C, Tamamaki N, Lovinger DM (2013): Voltage drives diverse endocannabinoid signals to mediate striatal microcircuit-specific plasticity. Nat Neurosci. 16: 1275–1283.

76. Gittis AH, Hang GB, LaDow ES, Shoenfeld LR, Atallah B V., Finkbeiner S, Kreitzer AC (2011): Rapid target-specific remodeling of fast-spiking inhibitory circuits after loss of dopamine. Neuron. 71: 858–868.

77. Voon V, Irvine MA, Derbyshire K, Worbe Y, Lange I, Abbott S, et al. (2014): Measuring “waiting” impulsivity in substance addictions and binge eating disorder in a novel analogue of rodent serial reaction time task. Biol Psychiatry. 75: 148–155.

