## Supplemental Data for "Nucleus Accumbens Fast-Spiking Interneurons Constrain Impulsive Action"

**SUPPLEMENTAL METHODS**

**5-Choice Serial Reaction Time Task (5-CSRTT)**

Each trial was signaled by the offset of a house light, followed by illumination of cue lights located inside five nose-poke apertures on the wall across from the food magazine. In Stage 1 of training, cues were immediately and continuously presented in all five apertures, and a response in any aperture was rewarded. In Stage 2 of training, a cue light was immediately and continuously presented in only one aperture. Correct responses in this aperture were rewarded, while incorrect responses in any other aperture were punished with illumination of the house light during a time-out period (five seconds). Mice moved on to the next stage after receiving 50 rewards before the end of each 60-minute session.

In Stage 3 and subsequent stages, each trial began with an inter-trial interval (ITI) prior to cue presentation. Mice were required to withhold responses during the ITI, and premature responses during this period were punished with a time-out. Trials were not re-set following a premature response. The duration of cue presentation was also limited, and failures to respond ("omissions") were punished with a time-out. Correct responses to the cued aperture were rewarded, and additional nose-pokes that followed a correct response but preceded retrieval of the reward were recorded as perseverative responses. Mice progressed to the next stage after reaching 50% accuracy, defined as the number of correct trials over the total number of all trial types. At various stages of training in different experiments, we also conducted tests of impulsivity and attention, in which either the ITI length or cue duration varied randomly from trial to trial (2.5-20 sec) within a single session. The impulsivity and attention tests ended after 100 trials (25 in each condition).

**Stereotaxic Surgery**

Mice were anesthetized with a ketamine:xylazine cocktail (100:10 mg/kg), and a small hole was drilled above target coordinates for the NAc core (AP +1.35, ML ±1.10; DV -4.40). A 33-gauge Hamilton syringe containing viral solution was slowly lowered to these target coordinates, followed by injection of 0.75 µL at a rate of 0.1 µL/min. The syringe tip was left in place at the injection site for 5 minutes, and then slowly retracted over the course of 5 minutes. For fiber photometry experiments, a unilateral 400 µm fiber-optic cannula (Doric Lenses: MFC_400/430-0.48_6mm_MF1.25_FLT) was implanted just above the site of virus injection (+0.02 mm), and fixed to the scull using a dual-cure resin (Patterson Dental, Inc.). For optogenetic experiments, bilateral 200 µm fiber optic implants (ThorLabs: 0.5NA fiber and ceramic ferrule) were positioned above the site of virus injection (+0.1 mm), and affixed similarly to fiber photometry experiments. Precautions were taken to prevent optogenetic light from influencing behavior by shielding patch cables/connections and applying opaque nail polish to headcaps.

**Fiber Photometry**

Real-time data were acquired using a RZ5P fiber photometry workstation (Tucker Davis Technologies). As previously described (1), 470 nm and 405 nm LEDs (ThorLabs) were modulated at distinct carrier frequencies (531 Hz and 211 Hz, respectively), and passed through a fluorescence mini cube (Doric Lenses) coupled into a patch cord (400 µm, 0.48 NA). For fiber photometry experiments involving red-shifted optogenetic excitation (ChrimsonR) or inhibition (eNpHR3.0), a 595nm LED (ThorLabs) was connected to the fluorescence mini cube and coupled to the same patch cord, and controlled using a pulse generator (Master-8, A.M.P.I.). The distal end of the patch cord was connected to the implanted fiberoptic cannula with a ceramic sleeve. Fluorescence was back-projected through the mini cube and focused onto a photoreceiver (Newport). Signals were sampled at 6.1 kHz, demodulated in real-time, and saved for offline analysis. For the 5-CSRTT, behaviorally relevant events (including trial onset, cue onset/offset, nose poke, and magazine entry) were transmitted to the fiber photometry workstation as TTL signals from the operant chambers. The change in fluorescent signal (dF/F) was calculated within each session as described (1). Each channel was low-passed filtered (<2Hz) and a linear least-squared model fit the isosbestic control signal (405 nm) to the calcium-dependent signal (470 nm). Change in fluorescence was calculated as ([470nm signal − fitted 405nm signal]/[fitted 405nm signal]). We also subtracted the eighth percentile value of dF/F over a rolling 20 sec window (2), which effectively corrected for photobleaching over time.

For event-related analyses, the photometry signal on each trial was first normalized to a 2 sec baseline period immediately preceding trial onset. Signals were then aligned to either trial onset, cue presentation, nose-poke response, or reward retrieval. It was necessary to align photometry data to each of these task events independently, because the temporal interval between events varied from trial to trial as function of the randomly selected ITI (5-10 sec) and the animal’s response latency. After aligning the fiber photometry signal to trial onset, we analyzed the average signal during the first 2 sec immediately following trial onset. After aligning the fiber photometry signal to cue presentation and nose-poke response, we analyzed the average signal during the 2 sec immediately preceding each event.

#### Fluorescent *in situ* Hybridization

PV-2A-Cre mice expressing AAVdj-hSyn-FLEX-ChrimsonR-tdTomato in the NAc were anesthetized with isoflurane and decapitated. Brains were rapidly frozen on dry ice in OCT embedding medium, and coronal sections (10 µm) were collected on positively charged glass slides and stored at -80°C until processing. Fluorescent *in situ* hybridization was performed using the RNAscope Multiplex Fluorescent Assay (Advanced Cell Diagnostics) according to manufacturer protocol. Briefly, the day before running the assay, slides were post-fixed in 4% PFA for 15 min at 4°C, followed by dehydration in increasing concentrations of ethanol (50%>70%>100%) for 5 min each at room temperature (RT). After the final 100% ethanol wash, slides were stored at -20°C in 100% ethanol overnight. The following day, slides were left to air dry and a hydrophobic barrier was painted around sections to be analyzed. Protease IV was applied to sections of interest for 30 min at RT, followed by two brief washes in 1x PBS. Hybridization probes for *Pthlh* (456521), *Pvalb* (421931-C2), and tdTomato (317041-C3) were applied to sections and incubated at 40°C for 2 hours, then washed twice in 1x RNAscope buffer for 2 minutes. Amplification was performed using AMP1-4 detection reagents, which were applied sequentially and incubated for 15-30 min each at 40°C, washing with 1X RNAscope wash buffer between each reagent. After the final wash, slides were shaken dry and mounting medium (Invitrogen Prolong Gold antifade reagent with DAPI) was applied before coverslips were placed on the slides. Images were taken using a Keyence BZX fluorescence microscope under a 40X objective.

#### Electrophysiology

PV-2A-Cre mice expressing AAV8-hSyn-FLEX-hM4Di-mCherry, AAVdj-EF1a-DIO-CHieF-Venus, or AAV5-EF1a-DIO-eNpHR3.0-mCh in the NAc were anesthetized with isoflurane and decapitated. Brains were quickly removed and placed in ice-cold cutting solution containing (in mM): 228 sucrose, 26 $NaHCO_3$, 11 glucose, 2.5 KCl, 1 $NaH_2PO_4$-$H_2O$, 7 $MgSO_4$-$7H_20$, 0.5 $CaCl_2$-$2H_2O$. Coronal slices (240 µm thick) containing the NAc were collected using a vibratome (Leica VT1000S) and allowed to recover in a submerged holding chamber with artificial cerebrospinal fluid (aCSF) containing (in mM): 119 NaCl, 26.2 $NaHCO_3$, 2.5 KCl, 1 $NaH_2PO_4$-$H_2O$, 11 glucose, 1.3 $MgSO_4$-$7H_2O$, 2.5 $CaCl_2$-$2H_2O$. Slices recovered in warm ACSF (33°C) for 10-15 min and then equilibrated to room temperature for at least one hour before use. Slices were transferred to a submerged recording chamber and continuously perfused with aCSF at a rate of 2 mL/min at room temperature. All solutions were continuously oxygenated (95% $O_2$/5% $CO_2$).

Whole-cell recordings from FSIs or MSNs were obtained under visual control using IR-DIC optics. FSIs were distinguished from putative MSNs by the presence or absence of fluorescence, respectively, using an Olympus BX51W1 microscope. For current clamp recordings, borosilicate glass electrodes (3-5 MΩ) were filled with (in mM): 120 K-Gluconate, 20 KCl, 10 HEPES, 0.2 EGTA, 2 $MgCl_2$, 4 ATP-Mg, 0.3 GTP-Na (pH 7.2-7.3). Cells were injected with a series of current steps (1 sec duration) from -100 to +250 pA. For voltage clamp recordings, cells were held at -70 mV using borosilicate glass electrodes were filled with (in mM): 120 $CsMeSO_4$, 15 CsCl, 10 TEA-Cl, 8 NaCl, 10 HEPES, 1 EGTA, 5 QX-314, 4 ATP-Mg, 0.3 GTP-Na (pH 7.2-7.3). For chemogenetic experiments, maximum firing rate and resting membrane potential were analyzed for each cell, before and after bath application of clozapine-N-oxide (CNO, 10 µM). Membrane potential measurements were corrected for a liquid junction potential of ~8 mV. Maximum firing rate was calculated as the average maximum firing rate over the 1 sec step that could be sustained without inducing a depolarization block. Action potential (AP) half-width was measured at the half-way point between threshold and the AP peak. For optogenetic stimulation of brain slices, ChIEF was activated at 475 nm (1-10 mW) and eNpHR3.0 was activated at 575 nm (~3 mW). Recordings were performed using a MultiClamp 700B (Molecular Devices), filtered at 2 kHz, and digitized at 10 kHz. Data acquisition and analysis were performed online using Axograph software. Series resistance was monitored continuously and experiments were discarded if resistance changed by >20%.

**Immunohistochemistry**

To confirm virus expression and/or fiber optic location, all mice were deeply anaesthetized using Beuthanasia (200 mg/kg, i.p.) and transcardially perfused with PBS followed by ice-cold 4% paraformaldehyde in PBS. Brains were fixed overnight in 4% paraformaldehyde in PBS then sliced at 40 µm thickness using a vibratome (Leica VT1000S). Free-floating coronal sections containing the NAc were incubated for 3hrs in blocking solution containing 0.2% Triton-X, 2% normal horse serum, and 0.05% Tween20. Sections were then incubated for 72 hrs with primary antibodies (mouse anti-mCherry 1:1000, Abcam; mouse anti-eYFP 1:1000,

Abcam; rabbit anti-RFP 1:1000, Rockland), washed five times, and incubated overnight with secondary antibodies (donkey anti-rabbit Alexa Fluor 647 IgG 1:1000, Life Technologies; goat anti-mouse Alexa Fluor 405, 488 or 647 IgG 1:1000, Abcam). Slides were washed and counter-stained for 20 minutes with 4',6-diamidino-2-phenylindole (DAPI) (1:50,000, Life Technologies), and mounted. Fluorescent images were acquired on a Keyence microscope using a 20x or 40x objective and viewed using ImageJ software.

**Open Field Behavior**

We tested open-field locomotor activity in mice having completed chemogenetic experiments using an arena (ENV-510, Med Associates) housed within a sound-attenuating chamber. The location of the mouse within the arena was tracked in two dimensions by arrays of infrared beams connected to a computer running Activity Monitor software (Med Associates). All mice were acclimated to the arena for one session on the day prior to testing, and then received i.p. injection of saline or CNO (2 mg/kg) on separate days in counterbalanced order, 30 minutes prior to a 60 minute test session.

**Statistics**

The absolute numbers of each trial type in the 5-CSRTT are presented in Supplementary Tables S2 and S3. Because there was no difference in the total number of completed trials between conditions, data for each trial type is graphically represented as a percentage of total trials. These data were further reduced to difference scores, allowing us to analyze statistical interactions between trial type and group, which provides a stringent test for the specificity of a given manipulation in experimental versus control groups (3).

Analysis of variance (ANOVA) was conducted in IBM SPSS Statistics v24, using either one-way or factorial models as appropriate, and with repeated measures on within-subject factors. Significant main effects and interactions were followed with LSD post-hoc tests. For main effects or interactions involving repeated measures, the Huynh-Feldt correction was applied to control for potential violations of the sphericity assumption. This correction reduces the degrees of freedom, resulting in non-integer values. The Type I error rate was set to $\alpha = 0.05$ (two-tailed) for all comparisons. Effect sizes are expressed as partial eta-squared ($\eta_p^2$) values, and used to compare effect magnitude in male and female mice because they are less influenced by unequal sample size than p values. All summary data are displayed as mean +/- SEM, with individual data points from male and female mice are shown as closed and open circles, respectively. Complete statistical results can be found in **Table S1**.

### SUPPLEMENTAL TABLES

***Supplemental Table S1: Comprehensive reporting of statistical results from ANOVA models.***

-included as an e-component in Excel format

***Supplemental Table S2. Absolute trial numbers for chemogenetic experiments in the 5-CSRTT (stage 8)***

| **Cre+** | | **Premature** | **Correct** | **Incorrect** | **Omission** | **Total** |
|---|---|---|---|---|---|---|
| Saline | Mean | 7.71 | 50.00 | 2.57 | 77.79 | 138.07 |
| | S.E. | 1.86 | 0.00 | 0.67 | 12.86 | 14.19 |
| CNO | Mean | 18.57 | 50.00 | 3.79 | 68.29 | 140.64 |
| | S.E. | 3.75 | 0.00 | 0.99 | 10.43 | 11.78 |
| *Statistical comparison between conditions (one-way ANOVA)* | | | | | | |
| | *F value* | *6.808* | *N/A* | *2.546* | *0.569* | *0.013* |
| | *p value* | *0.023* | *N/A* | *0.137* | *0.465* | *0.912* |
| | $\eta_p^2$ | *0.362* | *N/A* | *0.175* | *0.045* | *0.001* |
| **Cre-** | | **Premature** | **Correct** | **Incorrect** | **Omission** | **Total** |
| Saline | Mean | 10.43 | 50.00 | 2.57 | 44.07 | 107.07 |
| | S.E. | 1.79 | 0.00 | 0.69 | 9.57 | 10.76 |
| CNO | Mean | 9.93 | 50.00 | 2.86 | 32.71 | 95.36 |
| | S.E. | 2.22 | 0.00 | 0.88 | 5.45 | 5.52 |
| *Statistical comparison between conditions (one-way ANOVA)* | | | | | | |
| | *F value* | *0.049* | *N/A* | *0.238* | *1.104* | *0.939* |
| | *p value* | *0.828* | *N/A* | *0.635* | *0.314* | *0.352* |
| | $\eta_p^2$ | *0.004* | *N/A* | *0.019* | *0.084* | *0.073* |

***Supplemental Table S3. Absolute trial numbers for optogenetic experiments in the 5-CSRTT (stage 8)***

| **eNpHR3.0** | | **Premature** | **Correct** | **Incorrect** | **Omission** | **Total** |
|---|---|---|---|---|---|---|
| Control (Dark) | Mean | 1.89 | 22.78 | 1.00 | 26.44 | |
| | S.E. | 0.59 | 0.97 | 0.29 | 5.22 | |
| Cue Light | Mean | 1.22 | 20.67 | 3.00 | 27.22 | |
| | S.E. | 0.43 | 1.45 | 0.29 | 4.34 | |
| ITI Light | Mean | 10.11 | 16.56 | 0.44 | 24.67 | |
| | S.E. | 1.25 | 1.26 | 0.24 | 4.53 | |
| All trials | Mean | | | | | 156.00 |
| | S.E. | | | | | 13.39 |
| *Statistical comparison between conditions (one-way ANOVA)* | | | | | | |
| *F value* | | *49.893* | *3.785* | *19.197* | *0.778* | |
| *p value* | | *<0.001* | *0.049* | *<0.001* | *0.478* | |
| $\eta_p^2$ | | 0.88 | 0.35 | 0.73 | 0.10 | |
| **eYFP** | | **Premature** | **Correct** | **Incorrect** | **Omission** | **Total** |
| Control (Dark) | Mean | 2.50 | 19.83 | 1.17 | 25.50 | |
| | S.E. | 0.89 | 0.95 | 0.40 | 3.02 | |
| Cue Light | Mean | 1.50 | 20.50 | 1.67 | 25.00 | |
| | S.E. | 0.50 | 0.85 | 0.84 | 2.34 | |
| ITI Light | Mean | 3.17 | 19.67 | 1.00 | 25.17 | |
| | S.E. | 1.14 | 1.41 | 0.68 | 2.59 | |
| All Trials | Mean | | | | | 146.67 |
| | S.E. | | | | | 7.05 |
| *Statistical comparison between conditions (one-way ANOVA)* | | | | | | |
| *F value* | | *2.576* | *0.099* | *0.703* | *0.058* | |
| *p value* | | *0.137* | *0.907* | *0.523* | *0.944* | |
| $\eta_p^2$ | | *0.392* | 0.024 | 0.149 | 0.014 | |

### SUPPLEMENTARY FIGURES

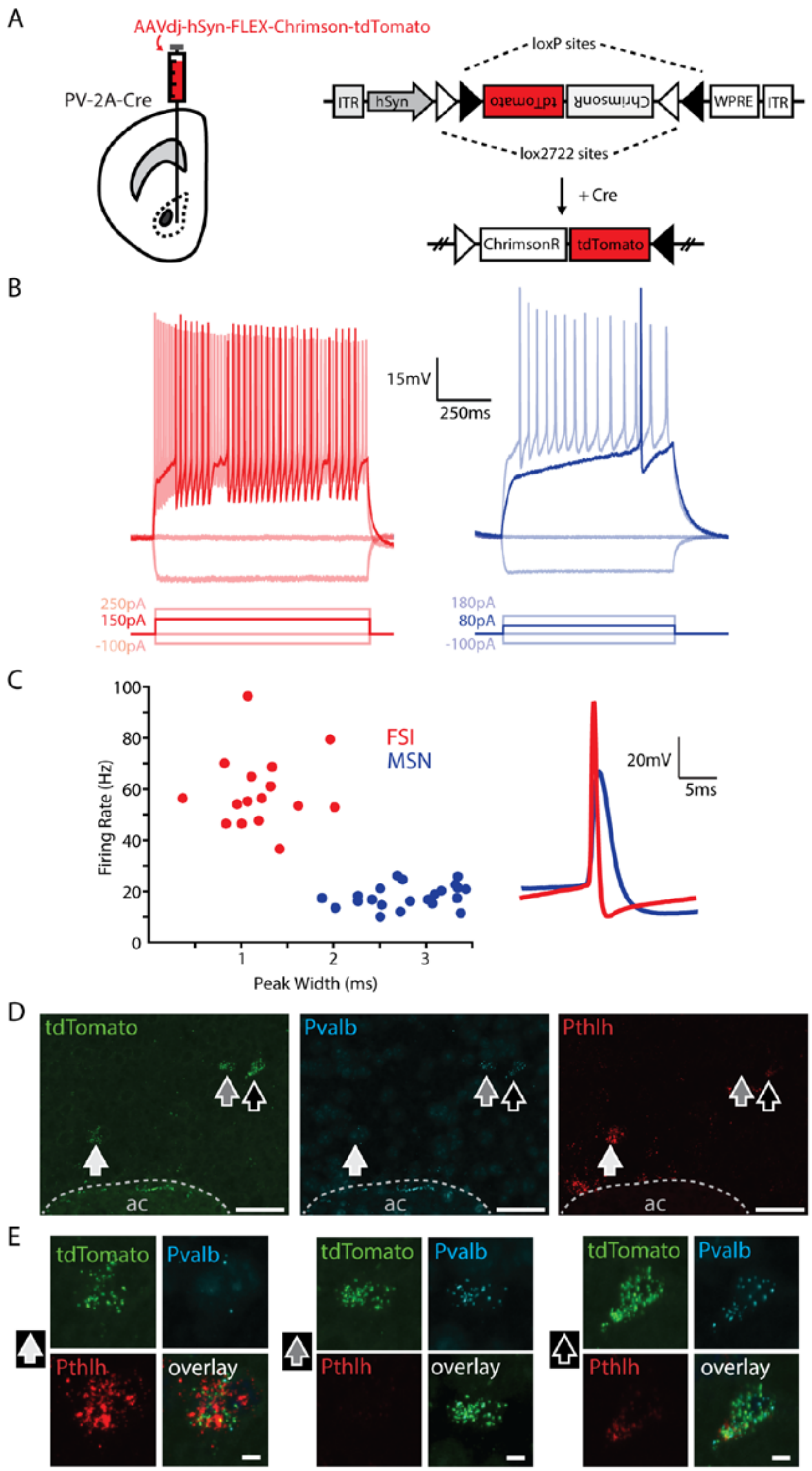

***Supplementary Figure S1. Electrophysiological and transcriptional characterization of virally-labeled FSIs in the NAc core**. (**A**) Viral strategy for labeling FSIs in PV-2A-Cre mice: flip-excision (FLEX) constructs are inverted and expressed by cells containing Cre recombinase. Note that bicistronic expression of ChrimsonR is not relevant for this experiment, but is used in other experiments. (**B**) Current-clamp recordings from (**left**) FSIs (red) and (**right**) medium spiny neurons (MSNs; blue) at regular current steps. Bolded step indicates lowest input current needed to elicit spiking (rheobase). (**C**) (**left**) Scatterplot of spike width at half amplitude (x) and maximum firing rate (y) for FSIs (red, n = 16) and MSNs (blue, n = 23). (**right**) Action potential waveforms of a representative FSI and MSN. (**D**) In situ hybridization of gene transcripts indicating co-expression of virally-expressed TdTomato (green) with Pvalb (cyan) and Pthlth (red) (scale bar, 50 µm; ac, anterior commissure). Arrows indicate cells in (E). (**E**) High magnification images showing Pvalb and Pthlh expression (scale bar, 5 µm).*

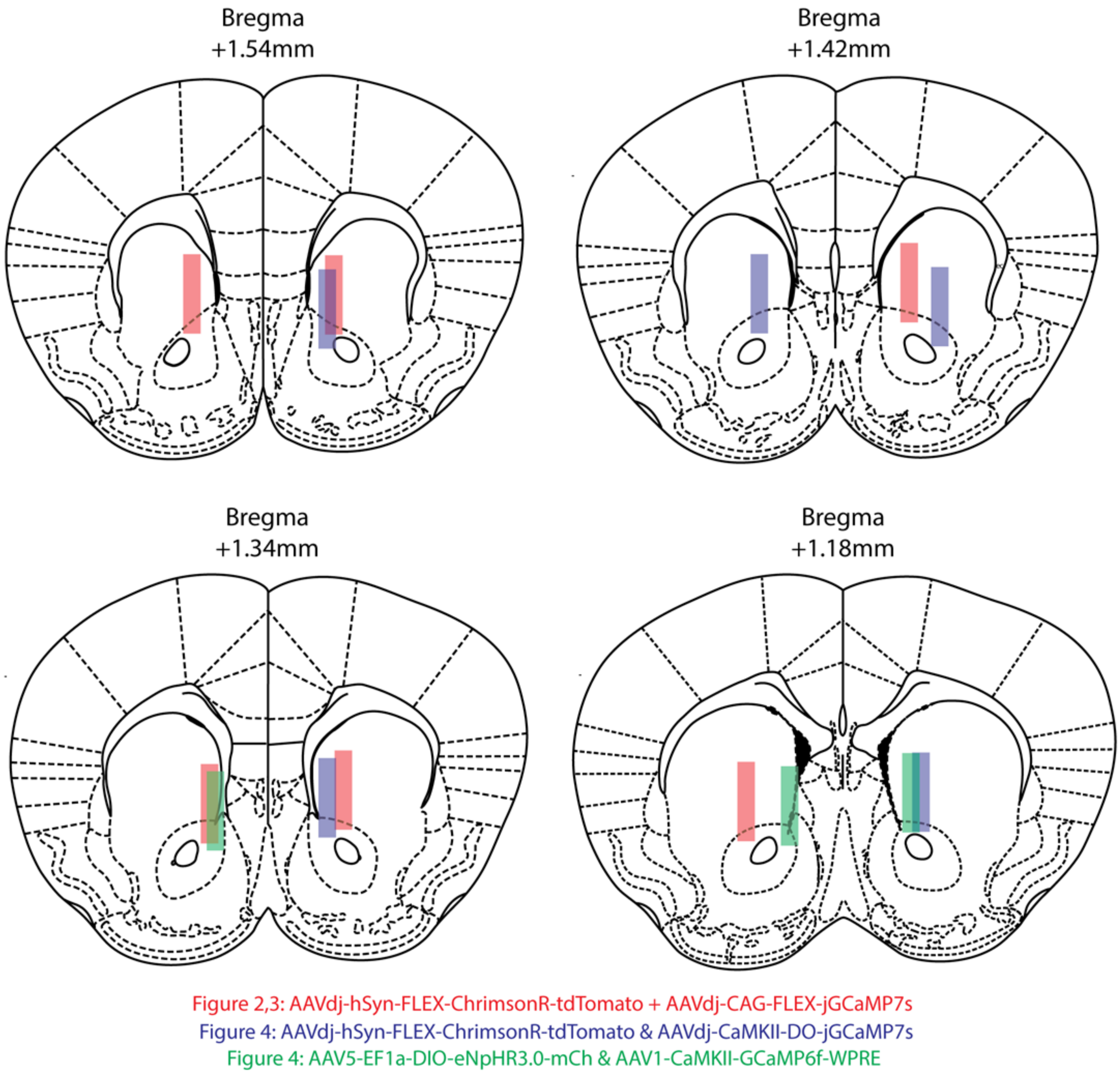

***Supplementary Figure S2. Anatomical localization of unilateral fiber photometry implants for experiments shown in Figures 2, 3, and 4.***

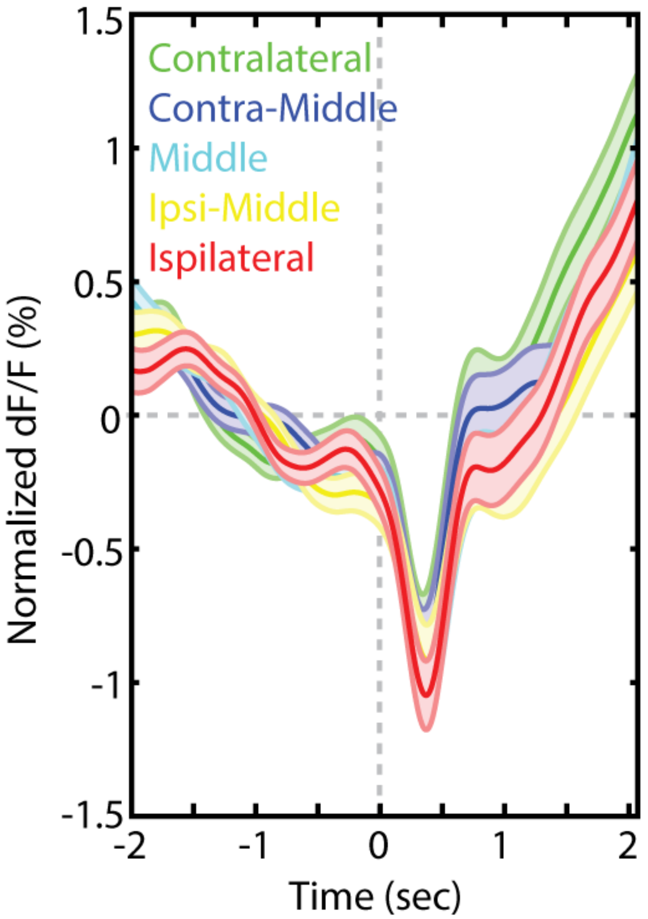

***Supplementary Figure S3. FSI activity measured with fiber photometry does not vary by nose-poke location.*** *Combined responses from PV-2A-Cre mice expressing jGCaMP7s on correct and premature trials in the 5-CSRTT are shown, with the location of nose poke response relative to the hemisphere in which fiber photometry signals were recorded. n=6 mice (3 female, 3 male).*

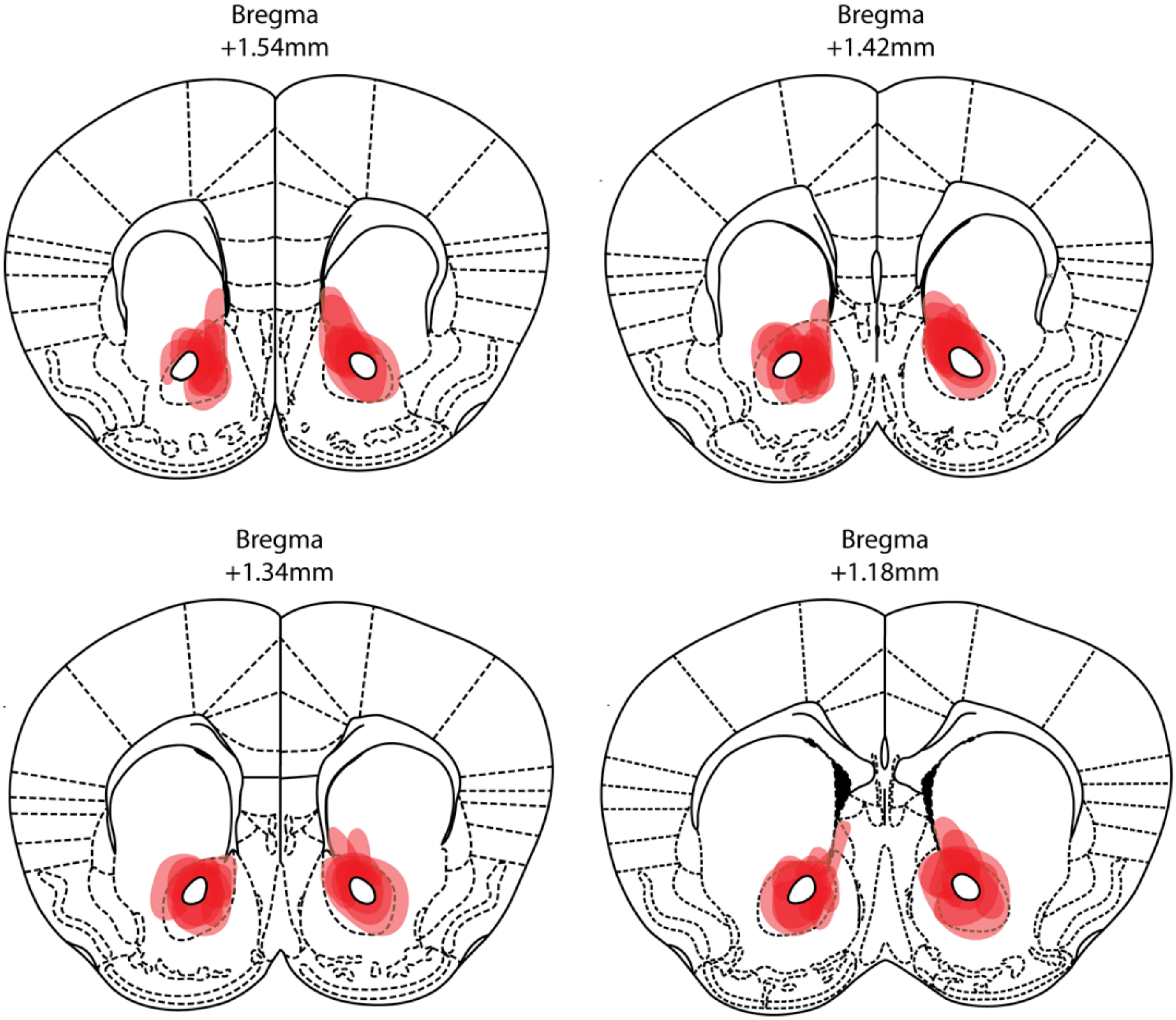

***Supplementary Figure S4. Anatomical localization of viral DREADD expression for chemogenetic experiments shown in Figure 5.***

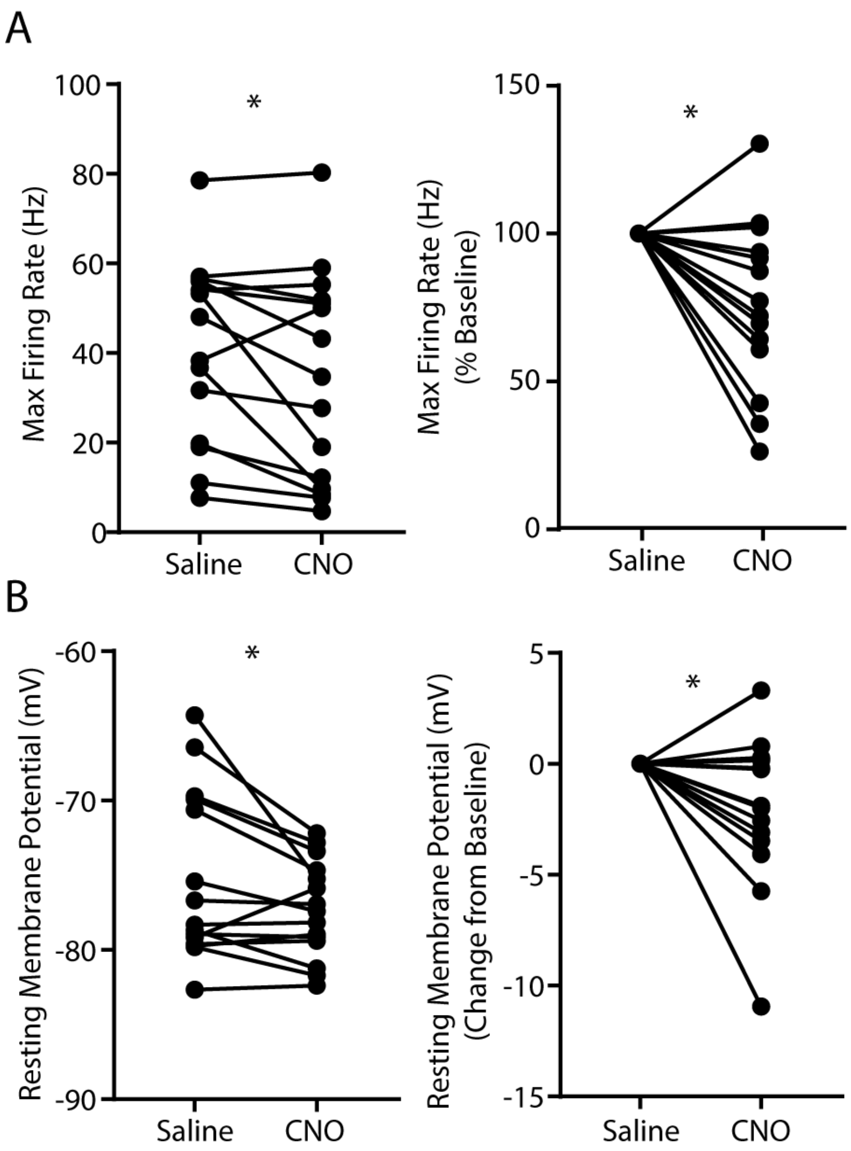

***Supplementary Figure S5. Electrophysiological validation for chemogenetic inhibition of FSIs.*** *Chemogenetic inhibition reduces (**A**) (**left**) raw and (**right**) normalized maximum spiking rate ($F_{1,14}$ = 5.73, p = 0.031,* $\eta_p^2$ = 0.29*) and (**B**) (**left**) raw and (**right**) normalized resting membrane potential ($F_{1,14}$ = 5.09, p = 0.041,* $\eta_p^2$ = 0.27*) of FSIs. n = 15 cells. *p<0.05.*

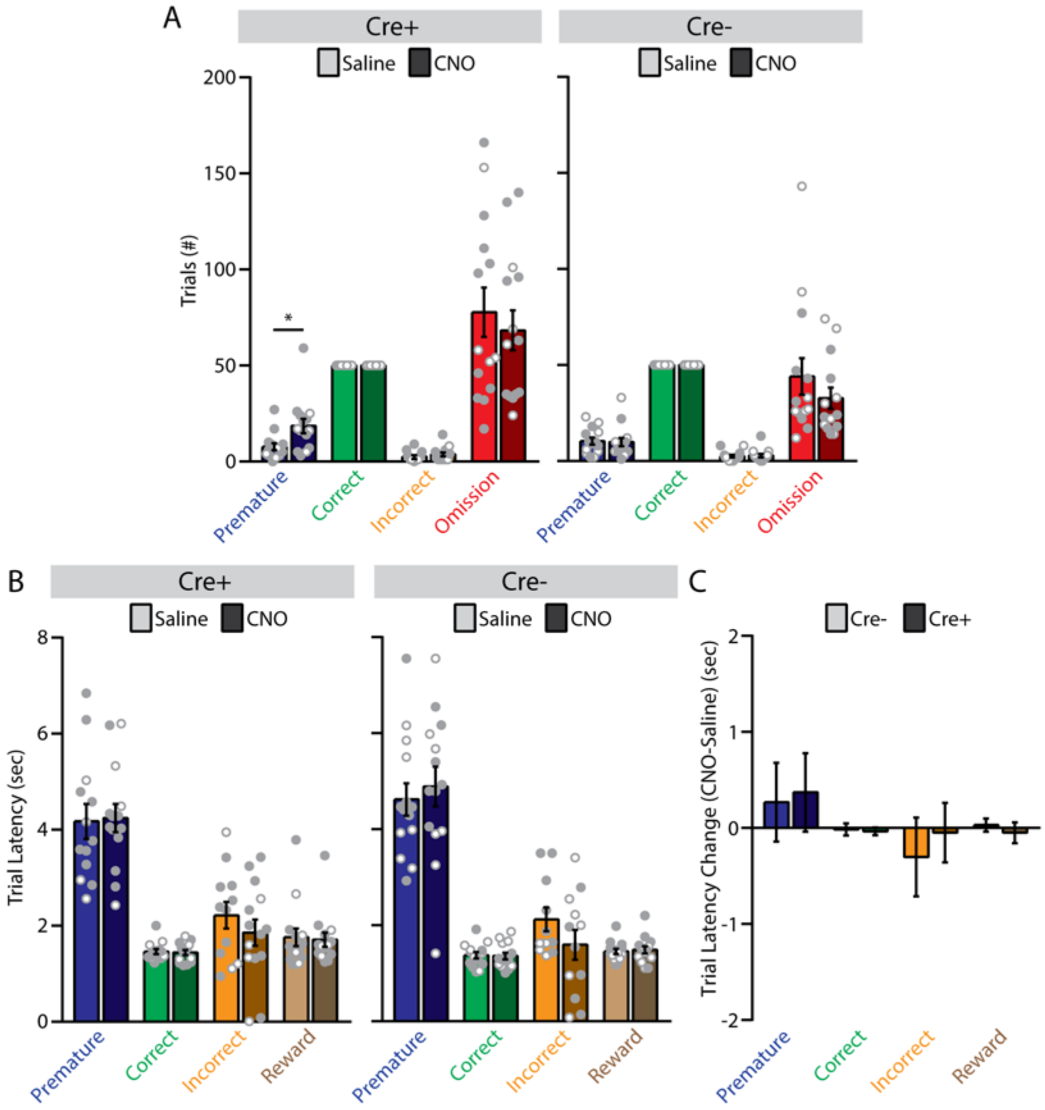

***Supplementary Figure S6. Additional data for chemogenetic experiments in the 5-CSRTT (stage 8). (A)** Trial numbers for (**left**) Cre+ and (**right**) Cre- mice. Cre+ mice showed a significant increase in number of premature trials compared to Cre- mice (Treatment: $F_{1,12}$ = 6.81, p = 0.023, $\eta_p^2$ = 0.36). **(B)** Latencies for (**left**) Cre+ and (**right**) Cre- mice. (**C**) Change in latencies (CNO – saline) for Cre- and Cre+ mice. *p<0.05.*

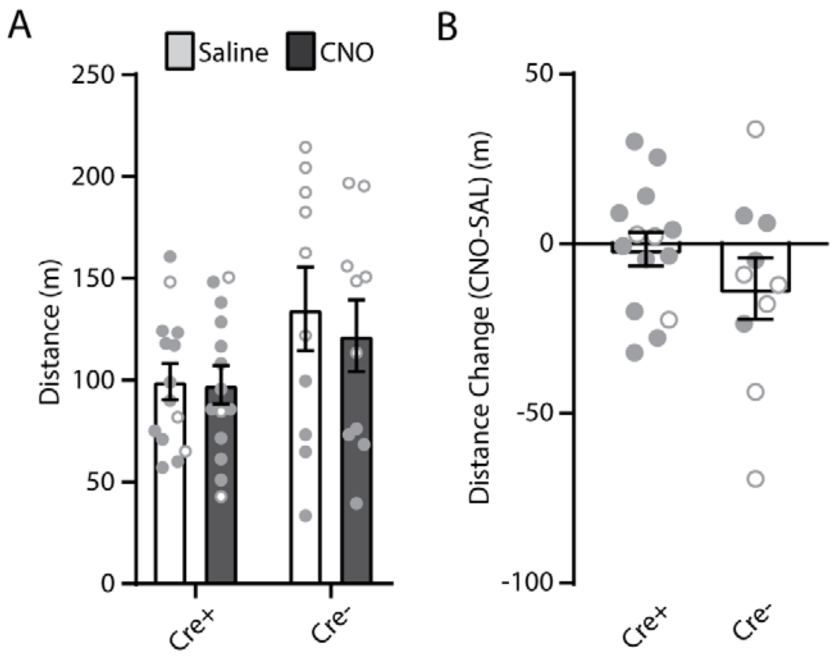

***Supplementary Figure S7. Chemogenetic inhibition of FSIs does not influence open-field locomotion. (A)*** *Raw distance traveled following saline or CNO (2 mg/kg) (Treatment x Genotype:* $F_{1,20} = 0.61$*,* $p = 0.445$*,* $\eta_p^2 = 0.03$*).* ***(B)*** *Change in distance traveled (CNO-saline) (Genotype:* $F_{1,20} = 2.09$*,* $p = 0.164$*,* $\eta_p^2 = 0.09$*). n = 14 Cre+ mice (11 male, 3 female) and 10 Cre- mice (4 male, 6 female).*

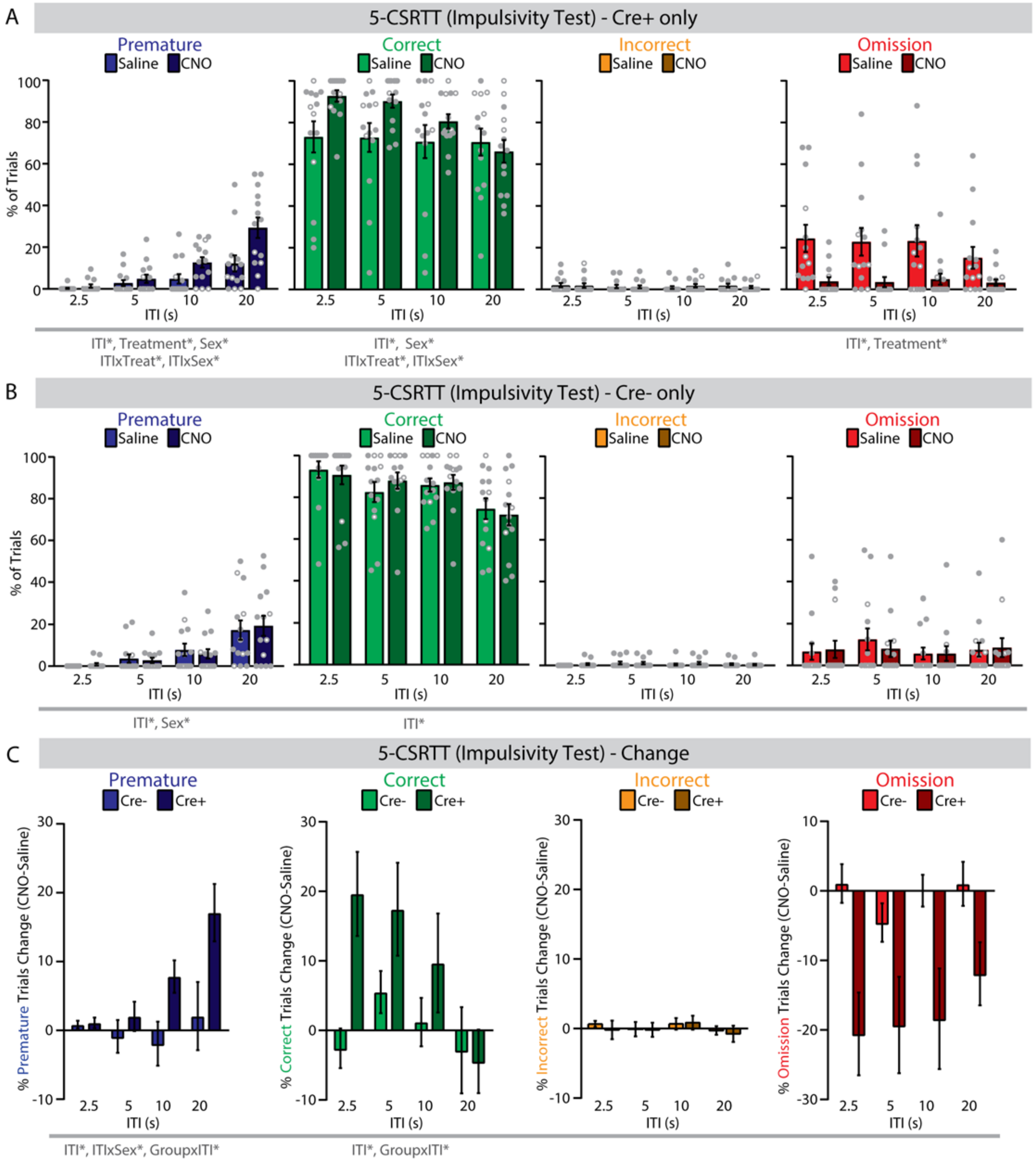

***Supplementary Figure S8. Additional data for chemogenetic experiments in the 5-CSRTT (impulsivity test)**. (**A**) In PV-2A-Cre mice, systemic administration of CNO (2 mg/kg) significantly increased the percentage of premature and correct trials, but did not change incorrect trials, and decreased omission trials). n = 14 mice; 10 male, 4 female. (**B**) In wild-type mice, systemic administration of CNO (2 mg/kg) did not change the percentage of premature, correct, incorrect, or omission trials. n = 14 mice; 7 male, 7 female. (**C**) There was no difference between PV-2A-Cre and wild-type mice in incorrect or omission trials, but a significant difference in correct and premature trials. *p<0.05. Full statistics can be found in Table S1.*

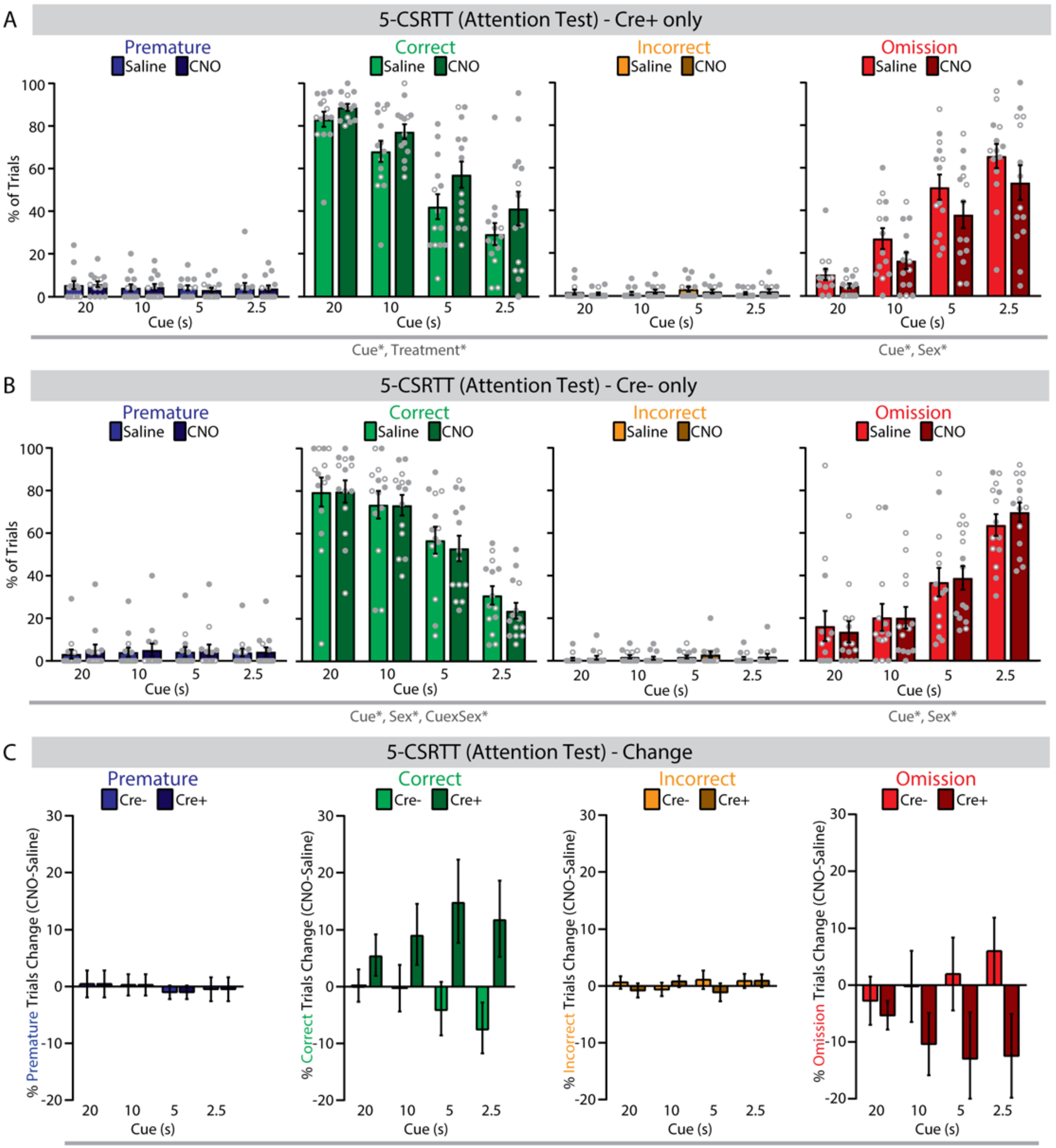

***Supplementary Figure S9. Additional data for chemogenetic experiments in the 5-CSRTT (attention test)**. (**A**) In PV-2A-Cre mice, systemic administration of CNO (2 mg/kg) significantly increased the percentage of correct trials). n = 14 mice; 10 male, 4 female. (**B**) In wild-type mice, systemic administration of CNO (2 mg/kg) did not change the percentage of premature, correct, incorrect, or omission trials. n = 14 mice; 7 male, 7 female. (**C**) There was no difference between PV-2A-Cre and wild-type mice in premature, correct, incorrect, or omission trials. *p<0.05. Full statistics can be found in Table S1.*

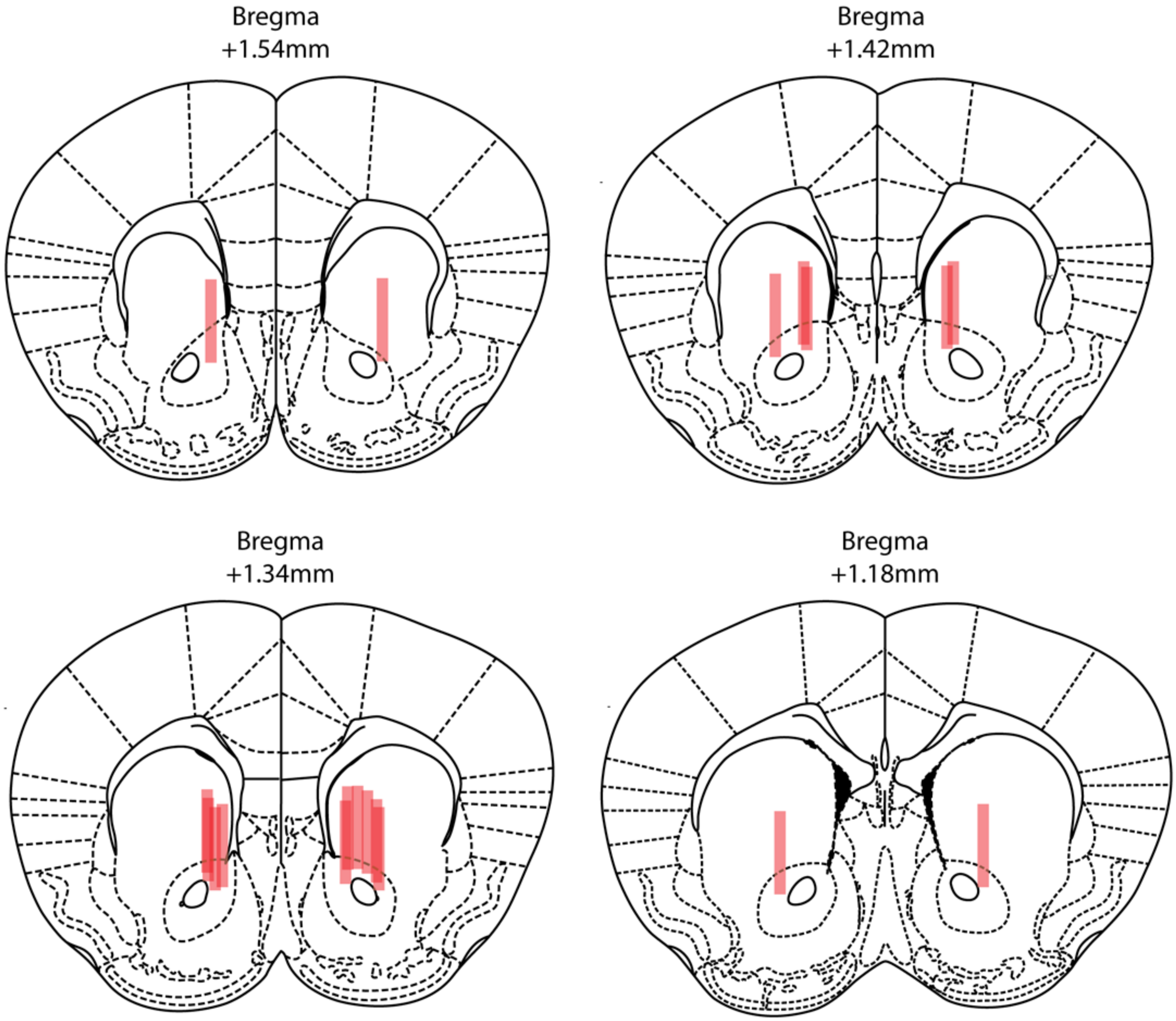

***Supplementary Figure S10. Anatomical localization of bilateral fiber-optic implants for optogenetic experiments shown in Figure 6.***

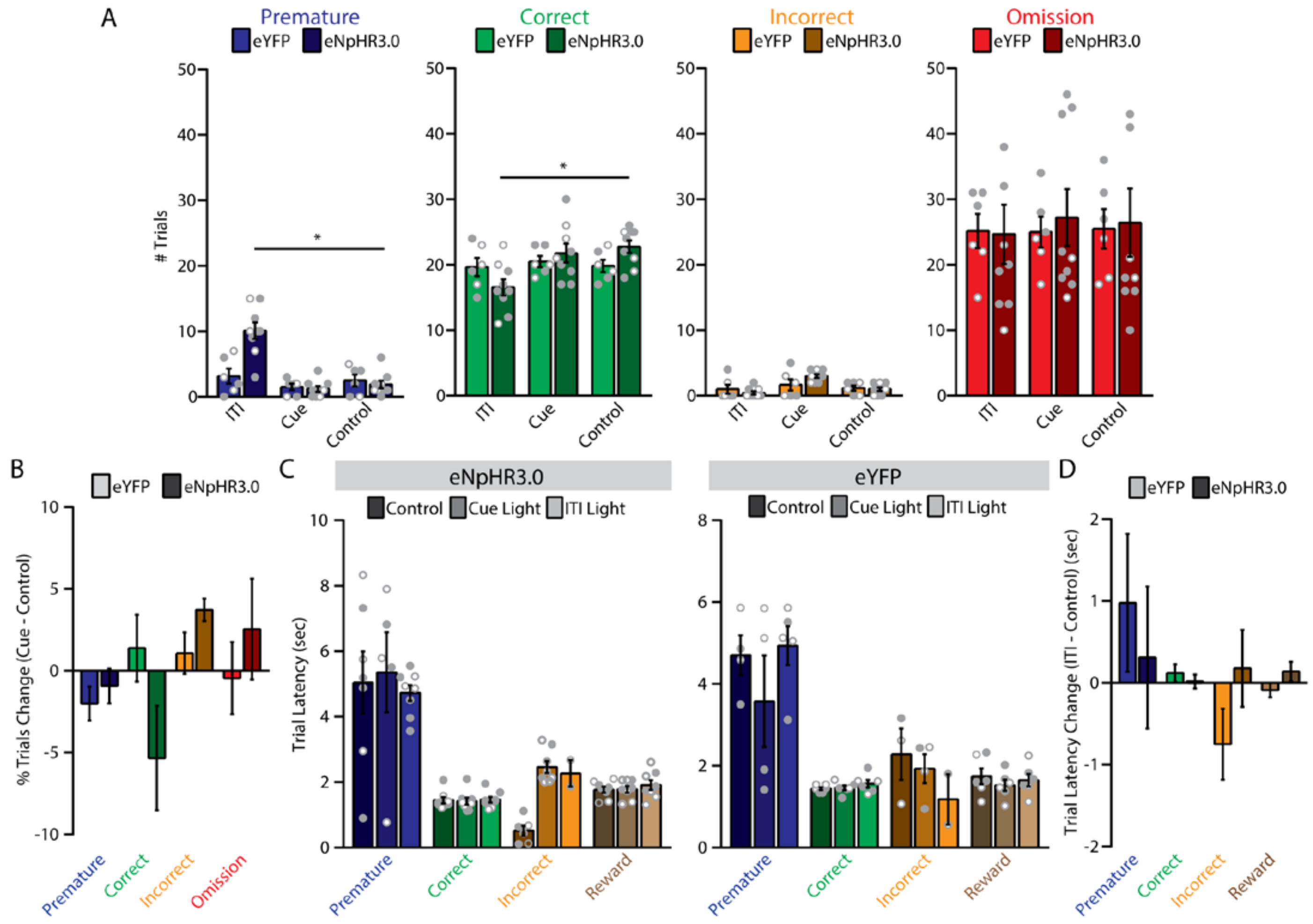

***Supplementary Figure S11. Additional data for optogenetic experiments in the 5-CSRTT (stage 8). (A)*** *Trial numbers involving light delivery during the ITI, during the cue, or no light (Control). Mice expressing eNpHR3.0 showed a significant increase in number of premature responses ($F_{2,14}$ = 49.89, p < 0.001, $\eta_p^2$ = 0.88) and decrease in number of correct responses ($F_{2,14}$ = 19.19, p < 0.001, $\eta_p^2$ = 0.73) with light delivery during the ITI. (**B**) Difference scores (Cue – Control) show no effect of light delivery during the cue. (**C**) Response latencies for trials involving light delivery during the ITI, during the cue, or no light (Control) for mice expressing (**left**) eNpHR3.0 or (**right**) eYFP. **(D)** Change in latencies (ITI - control) for mice expressing eNpHR3.0 or eYFP. n = 9 eNpHR3.0 mice (6 male, 3 female); 6 eYFP mice (3 male, 3 female). *p<0.05.*

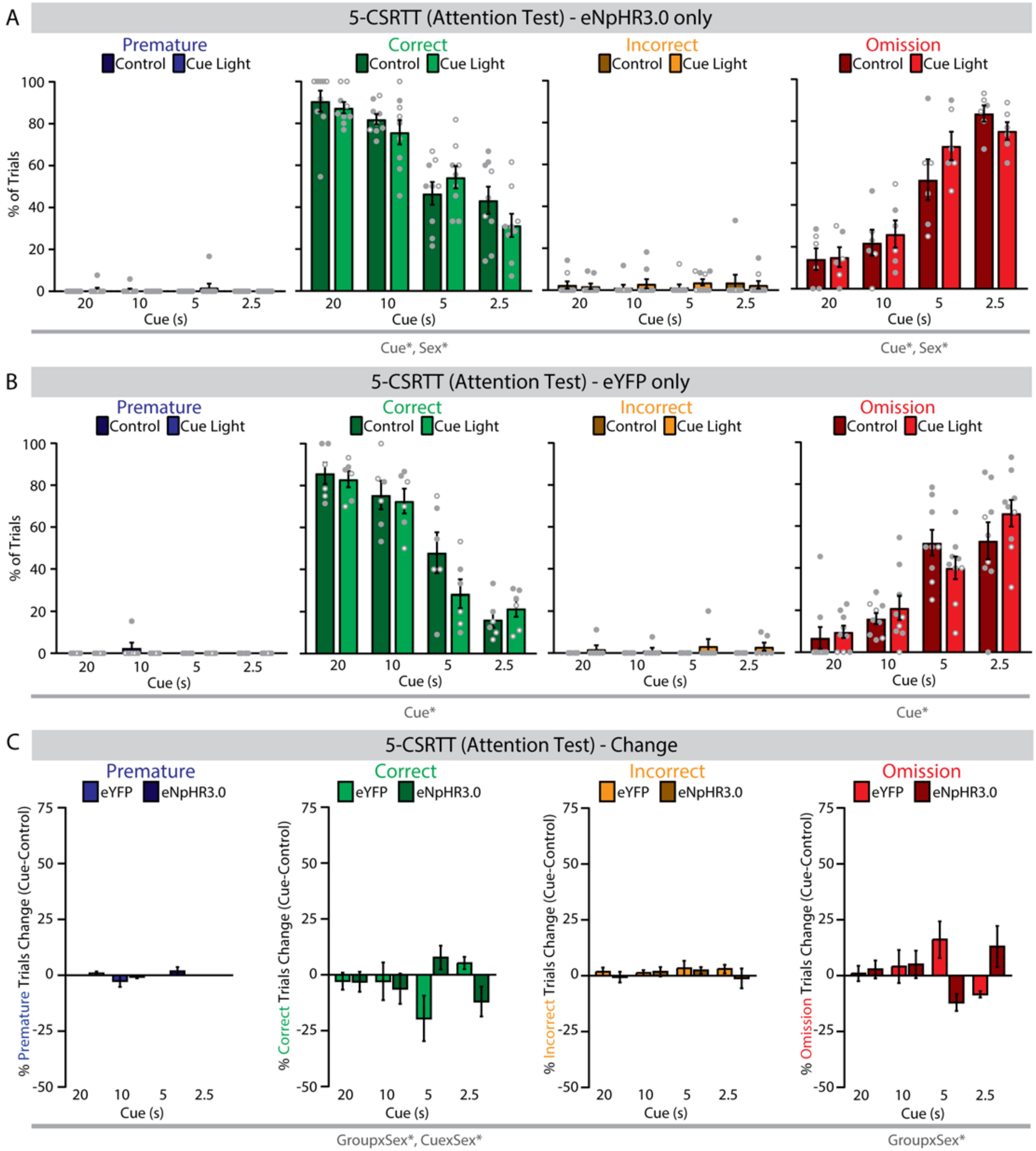

***Supplementary Figure S12. Additional data for optogenetic experiments in the 5-CSRTT (attention test)**. (**A**) In PV-2A-Cre mice expressing eNpHR3.0, light delivery during the cue had no effect on percentage of premature, correct, incorrect, or omission trials, compared to control (no light) trials. n = 9 mice; 6 male, 3 female. (**B**) In PV-2A-Cre mice expressing eYFP, light delivery during the cue had no effect on percentage of premature, correct, incorrect, or omission trials, compared to control (no light) trials. n = 6 mice; 3 male, 3 female. (**C**) There was no difference between PV-2A-Cre mice expressing eNpHR3.0 or eYFP in premature or incorrect trials, but a significant difference in correct and omission trials. *p<0.05. Full statistics can be found in Table S1.*

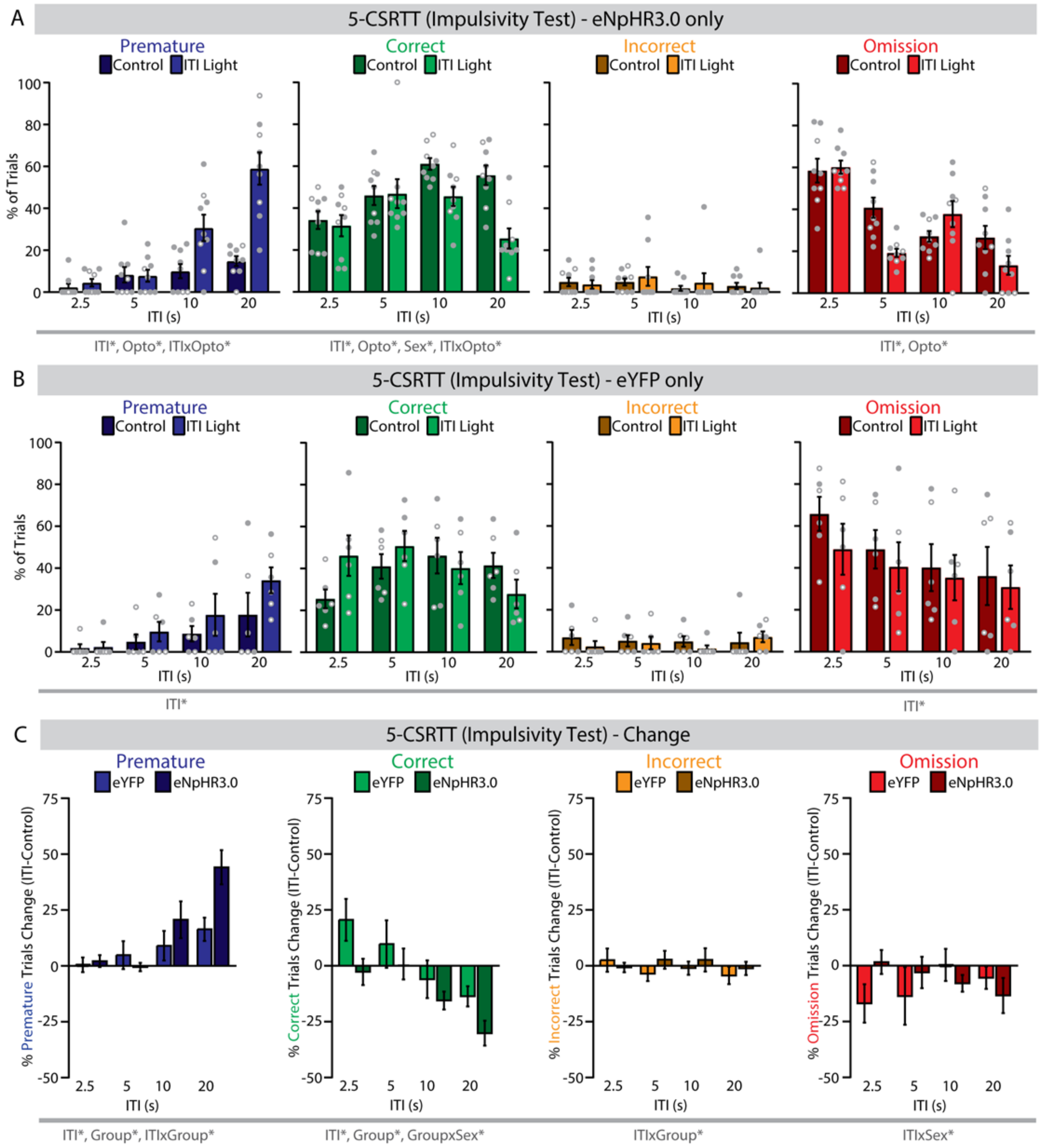

***Supplementary Figure S13. Additional data for optogenetic experiments in the 5-CSRTT (impulsivity test)***. *(**A**) In PV-2A-Cre mice expressing eNpHR3.0, light delivery during the inter-trial interval (ITI) significantly increased premature trials, decreased correct trials, and decreased omissions compared to control (no light) trials. n = 9 mice; 6 male, 3 female. (**B**) In PV-2A-Cre mice expressing eYFP, light delivery during the ITI had no effect on percentage of premature, correct, incorrect, or omission trials, compared to control (no light) trials. n = 6 mice; 3 male, 3 female. (**C**) There was no difference between PV-2A-Cre mice expressing eNpHR3.0 or eYFP in incorrect or omission trials, but a significant difference in premature and correct trials. *p<0.05. Full statistics can be found in Table S1.*

### SUPPLEMENTAL REFERENCES

1. Lerner TN, Shilyansky C, Davidson TJ, Evans KE, Beier KT, Zalocusky KA, *et al.* (2015): Intact-brain analyses reveal distinct information carried by SNc dopamine subcircuits. *Cell*. 162: 635–647.
2. Hung LW, Neuner S, Polepalli JS, Beier KT, Wright M, Walsh JJ, *et al.* (2017): Gating of social reward by oxytocin in the ventral tegmental area. 1411: 1406–1411.
3. Nieuwenhuis S, Forstmann BU, Wagenmakers E-J (2011): Erroneous analyses of interactions in neuroscience: a problem of significance. *Nat Neurosci*. 14: 1105–7.
